# Mesenchymal-epithelial transition serves to rapidly, yet transiently, restore the endometrial epithelium during postpartum murine uterine regeneration

**DOI:** 10.1101/2025.05.05.652287

**Authors:** Zidao Wang, Kimberly M. Davenport, Susanta K. Behura, Amanda L. Patterson

## Abstract

The uterus is a remarkable organ in its ability to undergo extensive tissue damage during menstruation and parturition, yet achieves efficient, scar-free repair. Coordinated regulation of this regenerative process is essential for uterine homeostasis and fertility; however, the underlying mechanisms remain incompletely understood. Here, we demonstrate that mesenchymal-epithelial transition (MET) contributed to postpartum endometrial re-epithelialization using *Pdgfrα^CreERT2/+^; Rosa26-tdTomato^fl/+^* lineage tracing mice. Flow cytometry revealed a marked increase in mesenchymal-derived (MD) epithelial cells during active tissue repair. Notably, these cells were transient, undergoing clearance primarily via apoptosis following completed epithelial restoration. We also identified a migratory population of transitional cells of mesenchymal origin within the mesometrial stroma that incorporated into the luminal epithelium, consistent with an active MET program. Single-nucleus RNA sequencing (snRNA-seq) revealed that MD epithelial cells exhibited gene expression profiles associated with cell adhesion and cytoskeletal remodeling, while transitional cells were enriched for genes involved in junctional assembly and actin dynamics. MET-associated genes were significantly upregulated in both transitional and MD epithelial populations. Cell-cell communication analysis highlighted WNT, BMP, and EPHA signaling as candidate regulators of MET during regeneration. Together, these findings provide confirmation of MET as a physiologic mechanism of postpartum endometrial epithelial repair and uncover a coordinated signaling network that facilitates this process. Perturbations in MET may contribute to pathologies such as endometriosis or endometrial cancer, underscoring the importance of understanding mesenchymal-epithelial plasticity in both normal and disease states.

**Significance:** The mammalian endometrium undergoes repeated injury and repair during menstruation (women) and pregnancy (most eutherians), yet exhibits a remarkable capacity for rapid, scar-free healing that mediates infection, inflammation and hemorrhage. Despite its clinical relevance, the molecular regulation of endometrial regeneration remains poorly defined. Using a transgenic lineage-tracing mouse model, we identified mesenchymal-derived (MD) epithelial and transitional cells during the regenerative window and revealed a critical role for mesenchymal-epithelial transition (MET) in this process. Single-nucleus RNA sequencing further uncovered functional characteristics of these cells and highlighted WNT, BMP, and EPHA signaling as potential regulators of MET. These findings provide new insight into the cellular and molecular framework of endometrial regeneration and have important implications for diseases involving aberrant tissue repair.

## Introduction

The adult uterus is a highly dynamic organ that undergoes coordinated structural and functional changes in response to cyclical fluctuations in ovarian hormones, mainly estrogen and progesterone. These cyclical changes, occurring as the menstrual cycle in women and the estrous cycle in other mammals, involve tightly regulated processes of cellular proliferation, differentiation, degeneration, and regeneration, with species-specific variations in the extent of tissue turnover. In women, approximately two-thirds of the endometrium (*i.e.,* the functional layer) is shed and subsequently regenerated from the basal layer nearly 400 times between puberty and menopause (1). In contrast, estrous-cycling species, such as mice, do not undergo overt endometrial shedding but exhibit hormone-driven cellular turnover through proliferation and apoptosis (2). In species with invasive embryo implantation, including humans and mice, the endometrium undergoes extensive remodeling during pregnancy and regenerates postpartum. Proper and timely endometrial regeneration is essential for preparing the uterus for subsequent reproductive cycles and pregnancy while preventing hemorrhage, mitigating hyperinflammation, protecting against infection, and maintaining tissue homeostasis.

Conversely, dysregulation of this process may lead to endometrial complications such as abnormal uterine bleeding (3), endometriosis (4), and endometrial cancer (5). Despite the extensive knowledge of the endocrine regulation of endometrial breakdown, the mechanisms underlying cellular regeneration of the endometrium are far from understood.

Restoration of the luminal epithelium (LE), termed re-epithelialization, is particularly crucial as it occurs during active endometrial dissolution and serves as a vital barrier to protect against infection. Several mechanisms have been proposed to facilitate endometrial re-epithelialization, including 1) replication of residual LE and glandular epithelium (GE) that escaped desquamation, 2) epithelial stem/progenitor cell activity, 3) infiltration and contribution of bone marrow-derived cells, and 4) mesenchymal-epithelial transition (MET) of endometrial stromal-mesenchymal cells (6).

MET is the process by which mesenchymal cells undergo a phenotypic and functional transformation into epithelial cells. Mesenchymal cells are spindle-shaped and highly migratory, but upon MET, they lose their migratory capacity, adhere to a basement membrane, and develop apical-basal polarity. Additionally, epithelial cells form cell-cell contacts by expressing various adhesion and junctional proteins such as epithelial cell adhesion molecule (EPCAM) and E-cadherin (CDH1). Both MET and the reverse process, epithelial-mesenchymal transition (EMT), play key roles in embryonic tissue and organ formation and wound healing in the adult but can contribute to pathologies, including fibrosis and cancer (7–9).

Accumulating literature provides evidence of MET occurring during endometrial re-epithelialization in the estrous cycle (10), following parturition (11, 12) and in menses-like regeneration (11, 13–15) in mouse models. In two foundational studies, mesenchymal lineage-tracing was performed using Anti-Müllerian Hormone Receptor Type 2 (*Amhr2)-Cre* mice, which drove the expression of either EYFP (11) or *LacZ* (12) reporters. EYFP-positive or β-gal-positive mesenchymal-derived (MD) epithelial cells were identified in the LE and GE following parturition and endometrial regeneration, suggesting the occurrence of MET. Further, by using the menses-like mouse model, migration of pan-cytokeratin and vimentin co-expressing “transitional” cells was tracked from the stromal-myometrial border (a proposed location of stromal stem/progenitor cells) to the mesometrial zone of regeneration (11). Subsequently, CD34^+^KLF4^+^ stromal cells were shown to contribute to the epithelium during menses-like endometrial regeneration, suggesting that a specific stromal stem/progenitor population served as the source of cells undergoing MET (14). Molecular evidence of cellular transition was demonstrated through the expression of EMT/MET-related genes throughout menses-like endometrial regeneration (13) and more recently single-cell RNA-sequencing (scRNA-seq) was used to investigate cells undergoing transition (15). Despite the growing body of literature in support of MET during endometrial re-epithelialization, this mechanism has also been called into question (16), substantiating the need for further investigation.

It is likely that MET occurs in women after parturition and during menstruation, a hypothesis first proposed as early as 1897 (17), although studies on this topic remain limited. In support of this, characterization of the hysteroscopic appearance of the endometrium during menstruation in women, revealed patches of nascent epithelial cells that were phenotypically distinct from the remnant mature GE cells. It was postulated that these cells originated from the differentiation of adjacent stromal cells, instead of the proliferation of residual glandular stumps (18, 19).

Despite our growing understanding of MET as a mechanism for endometrial re-epithelialization, knowledge gaps remain regarding the molecular regulation of MET and whether MD epithelial cells differ from resident non-MD epithelial cells. In the current study, we utilized an inducible lineage-tracing transgenic mouse model (*Pdgfrα^CreERT2/+^; Rosa26-tdTomato^fl/+^*) to investigate the role of endometrial stromal-mesenchymal cells in re-epithelialization via MET during postpartum endometrial regeneration. Through snRNA-seq, we provide the first evidence of the unique characteristics and functions of MD epithelial cells compared to resident non-MD epithelial cells. This study not only confirms the involvement of MET during endometrial re-epithelialization but also provides insights into an orchestrated signaling network underlying this process.

## Results

### Characterization of endometrial MD epithelial cells during endometrial regeneration

In our previous investigation of MET using *Amhr2^Cre/+^; Rosa26-EYFP^fl/+^*(Cre activated embryonically) mice it was discovered that MET may occur during postnatal uterine maturation and the estrous cycle, making it difficult to study MET specifically during regeneration (10).

Therefore, to control the timing of Cre recombination we adopted a tamoxifen (TAM)-inducible, lineage-tracing transgenic mouse model (*Pdgfrα^CreERT2/+^; Rosa26-tdTomato^fl/+^*) to investigate the role of endometrial stromal-mesenchymal cells in re-epithelialization via MET during postpartum endometrial regeneration (*SI Appendix,* **Fig. S1A**). Transgenic female mice were administered TAM (or vehicle) at sexual maturity to induce tdTomato (tdTom) expression in endometrial mesenchymal cells. Following a four-week TAM washout, uteri were collected from: 1) virgin females in estrus or diestrus, 2) at embryonic day (E) 18.5 before parturition and endometrial regeneration had begun, 3) postpartum days (PPD) 1-4 during endometrial regeneration, which was completed by PPD 4, and 4) after completed regeneration and resumption of estrous cyclicity at PPD28 (*SI Appendix,* **Fig. S1B**). The labeling efficiency of stromal cells in virgin mice by flow cytometry analysis (EpCAM^-^tdTom^+^/EpCAM^-^) was an average of 50.93% at estrus and 45.68% at diestrus (*SI Appendix,* **Fig. S1D and E**). Basal MET during the estrous cycle was assessed by quantifying the percentage of MD epithelial cells (EpCAM^+^tdTom^+^/EpCAM^+^) by flow cytometry, which averaged 0.915% in estrus and 2.098% in diestrus (*SI Appendix,* **Fig. S1D and F**). We also confirmed the native expression of PDGFRα protein by immunofluorescence showing co-localization with tdTom (*SI Appendix,* **Fig. S1G**).

Since we observed little MET during the estrous cycle, we proceeded to investigate the contribution of MET during postpartum endometrial re-epithelialization. With E18.5 to serve as the pre-endometrial regeneration time point, MD epithelial cells were present (0.71%) but comparable to that seen in cyclic mice (1.63%; P=0.7213; **Fig. 1B**). By contrast, a sharp increase in MD epithelial cells was observed from E18.5 (0.71%) to PPD1 (5.10%; P < 0.0001; **Fig. 1A and B**). MD epithelial cells peaked at PPD2 (6.92 %), followed by a steady decline until PPD4 (3.05%) and reached the basal level by PPD28 (1.73%; **Fig. 1A and B**). The presence of MD epithelial cells during endometrial regeneration was confirmed by immunofluorescent analysis of uterine tissue sections. Using an antibody to RFP, the tdTom signal was enhanced showing RFP^+^ cells present in both LE and GE interspersed with RFP^-^, non-MD (*i.e.,* resident) epithelial cells (**Fig. 1C**). These data indicate that MET occurs during the first 2 days of postpartum endometrial re-epithelialization and the resulting MD epithelial cells are rapidly removed as regeneration is completed.

**Figure 1.**
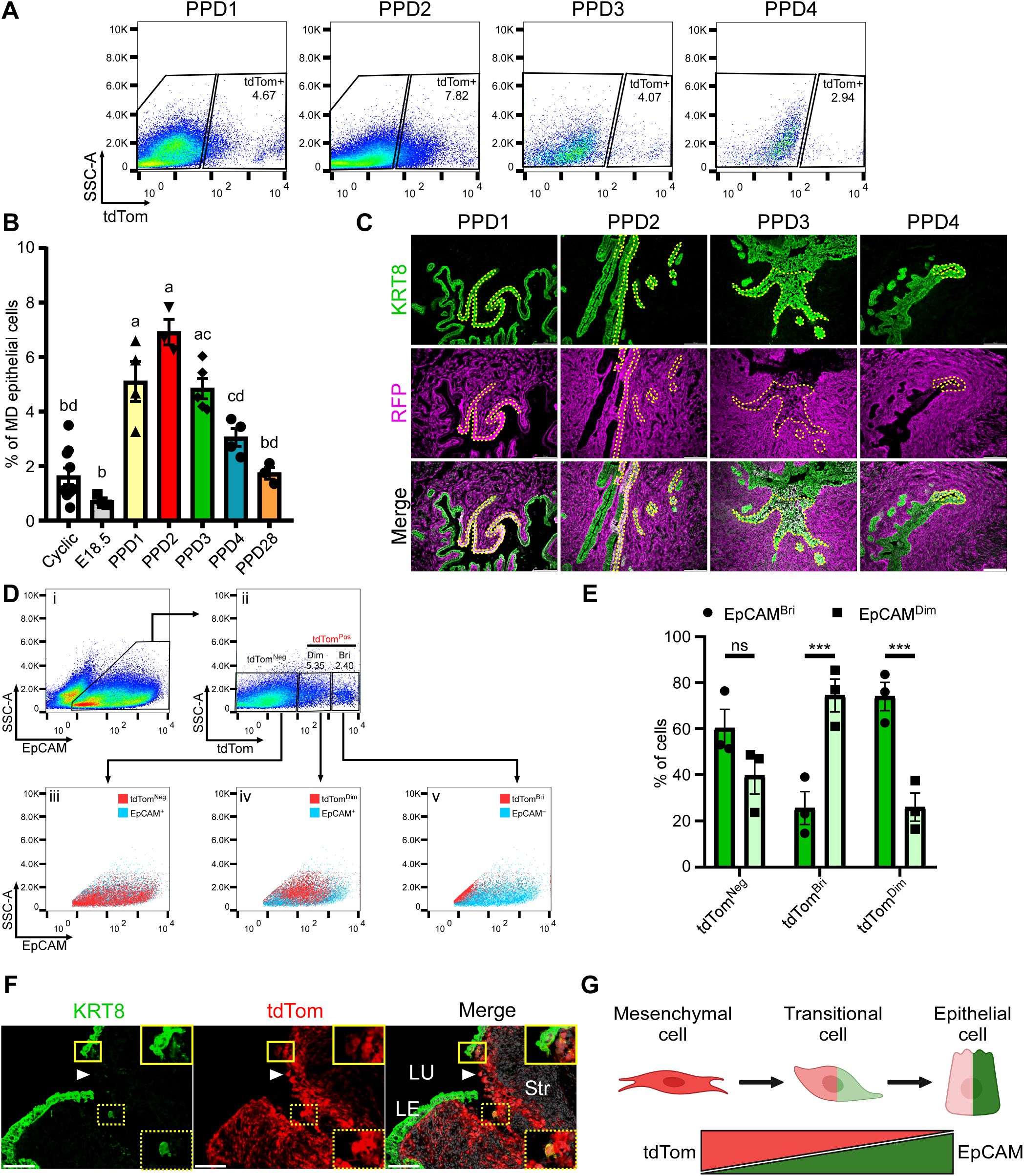
Mesenchymal-derived (MD) epithelial cells contribute to postpartum endometrial re-epithelialization. **(A)** Flow cytometry scatter plots showing tdTomato Red (tdTom) expression in the EpCAM^+^ epithelial population from postpartum day (PPD) 1 to PPD4 in uteri of *Pdgfrα^CreERT2/+^; Rosa26-tdTomato^fl/+^* mice. **(B)** Quantification of the percentage of EpCAM^+^tdTom^+^ MD epithelial cells in the uterine epithelia of cyclic and postpartum mice by flow cytometry analysis. **(C)** Immunofluorescence staining for KRT8 (green, top panel) and RFP (magenta, to amplify the tdTom signal, middle panel) with nuclear DAPI staining (grey, Merge, bottom panel) in uteri from PPD1 to PPD4. The dashed yellow lines outline regions of MD epithelial cells co-expressing KRT8 and RFP. **(D)** Flow cytometry scatter plots showing i) EpCAM expression; ii) tdTom expression in EpCAM^+^ epithelial cells subset as tdTom negative (tdTom^Neg^) and tdTom positive (tdTom^Pos^), which were further subset into tdTom dim (tdTom^Dim^) and tdTom bright (tdTom^Bri^); and backgating of iii) tdTom^Neg^, iv) tdTom^Dim^, and v) tdTom^Bri^ cells onto EpCAM expression. **(E)** Quantification of flow cytometry analyses of the percentage of EpCAM^Bri^ and EpCAM^Dim^ cells in each tdTom subset. **(F)** Immunofluorescence staining for KRT8 (green), with endogenous tdTom (red) and nuclear counterstaining with DAPI (grey, Merge) of a regeneration area in the uterus on PPD1. The dashed yellow box denotes a cell co-expressing dim KRT8 and bright tdTom, with an enlarged image inset. The solid yellow box denotes a patch of intact luminal epithelium (LE) with bright KRT8 and dim tdTom expression, with an enlarged image inset. The white arrowhead shows a stromal cell with no KRT8 expression at the denuded Lumen (LU). Str: stroma. Scale bar: 100 µm. **(G)** Schematic diagram illustrating reduction of tdTom labeling and induction of epithelial labeling during MET. Groups with different letters in (B) are statistically different (P < 0.05). ***P<0.001.

Amongst MD epithelial cells, we observed varied levels of tdTom expression by flow cytometry, allowing for segregation into tdTom dim (EpCAM^+^tdTom^Dim^) and tdTom bright (EpCAM^+^tdTom^Bri^) cells (**Fig. 1D-ii**). Back-gating of these two populations on EpCAM expression revealed that EpCAM^+^tdTom^Dim^ cells were mostly EpCAM bright (EpCAM^Bri^; **Fig. 1D-iv and E**), while EpCAM^+^tdTom^Bri^ cells were mostly EpCAM dim (EpCAM^Dim^; **Fig. 1D-v and E**), suggesting an inverse relationship between expression of epithelial markers and tdTom. This was visualized by immunofluorescence expression of the epithelial marker cytokeratin 8 (KRT8) in conjunction with tdTom expression in tissue sections of regenerating uteri. Cells with bright tdTom tended to have dimmer KRT8 and were un-incorporated into the LE (**Fig. 1F** dashed yellow inset). Conversely, cells with dim tdTom had brighter KRT8 and were more incorporated into the LE (**Fig. 1F**, solid yellow inset). During MET, mesenchymal cells lose mesenchymal properties while gaining epithelial properties. These data demonstrate the acquisition of epithelial markers suggesting active MET (**Fig. 1G**). Interestingly, tdTom expression appeared to decrease with completion of MET even though expression is controlled by the CAG promoter, not a mesenchyme-specific promoter. Literature suggests that CAG promoter activity can be impacted by the developmental state of the cells in which it is inserted (20, 21). Accordingly, diminished tdTom expression in MD epithelial cells may indirectly demonstrate a more differentiated state of these cells having undergone MET.

### *snRNA-seq of postpartum regenerating endometrium* reveals a unique transcriptomic profile of MD epithelial cells

Studies using different lineage-tracing mouse models have provided evidence of MET during postpartum and menses-like endometrial re-epithelialization (10–15). However, it is not fully understood if MD epithelial cells serve as *bona fide* epithelial cells that are indistinguishable from resident non-MD endometrial epithelial cells. Considering that MD epithelial cells arise and demise quickly during endometrial regeneration (**Fig. 1**), indicating their transient nature, it is likely that they have an altered phenotype compared to non-MD epithelial cells. To understand the transcriptomic profiles of MD epithelial cells, we performed snRNA-seq on postpartum uteri at all time points. After quality control, a total of 98,343 nuclei remained for downstream analyses (*SI Appendix,* **Fig. S2**). We first characterized known major cell types including mesenchymal, epithelial, endothelial, perivascular, myometrial, and immune cells **(Fig. 2A**). Of note, a distinct cluster expressing mesenchymal and epithelial markers was identified and designated as transitional cells (**Fig. 2A**, cluster 4, discussed below). The epithelial clusters were then subset by tdTom expression yielding tdTom^+^ MD and tdTom^-^ non-MD epithelial populations (**Fig. 2A**). To characterize the broad transcriptomic signature of MD epithelial cells, we compared the differentially expressed genes (DEGs) of non-MD and MD epithelial cells, mesenchymal, perivascular and myometrial cells (**Fig. 2B**). The top 10 DEGs of non-MD and MD epithelial cells included many genes commonly expressed in epithelial cells (*e.g Mecom, Alcam, Fut9, and Prom1)*, revealing a similar global transcriptomic profile of non-MD and MD epithelial cells compared to other major cell types. To reveal the transcriptomic features that distinguish MD from non-MD epithelial cells, DEG analysis was performed for the two populations, revealing 577 genes in total, 389 up- and 188 down-regulated (**Fig. 2C**, *SI Appendix* **Dataset S1**). Of interest were *Ceacam1* and *Trpv6* which function in cell junction organization and maintenance of epithelial cell integrity, respectively, and both are associated with cellular transitions. Deletion of *Ceacam1,* a cell adhesion molecule, in *Pten^+/-^* mice caused prostate intraepithelial neoplasia due to an increase in cell proliferation and epithelial-mesenchymal transition (EMT), suggesting an inhibitory effect of CEACAM1 on EMT (22).

**Figure 2.**
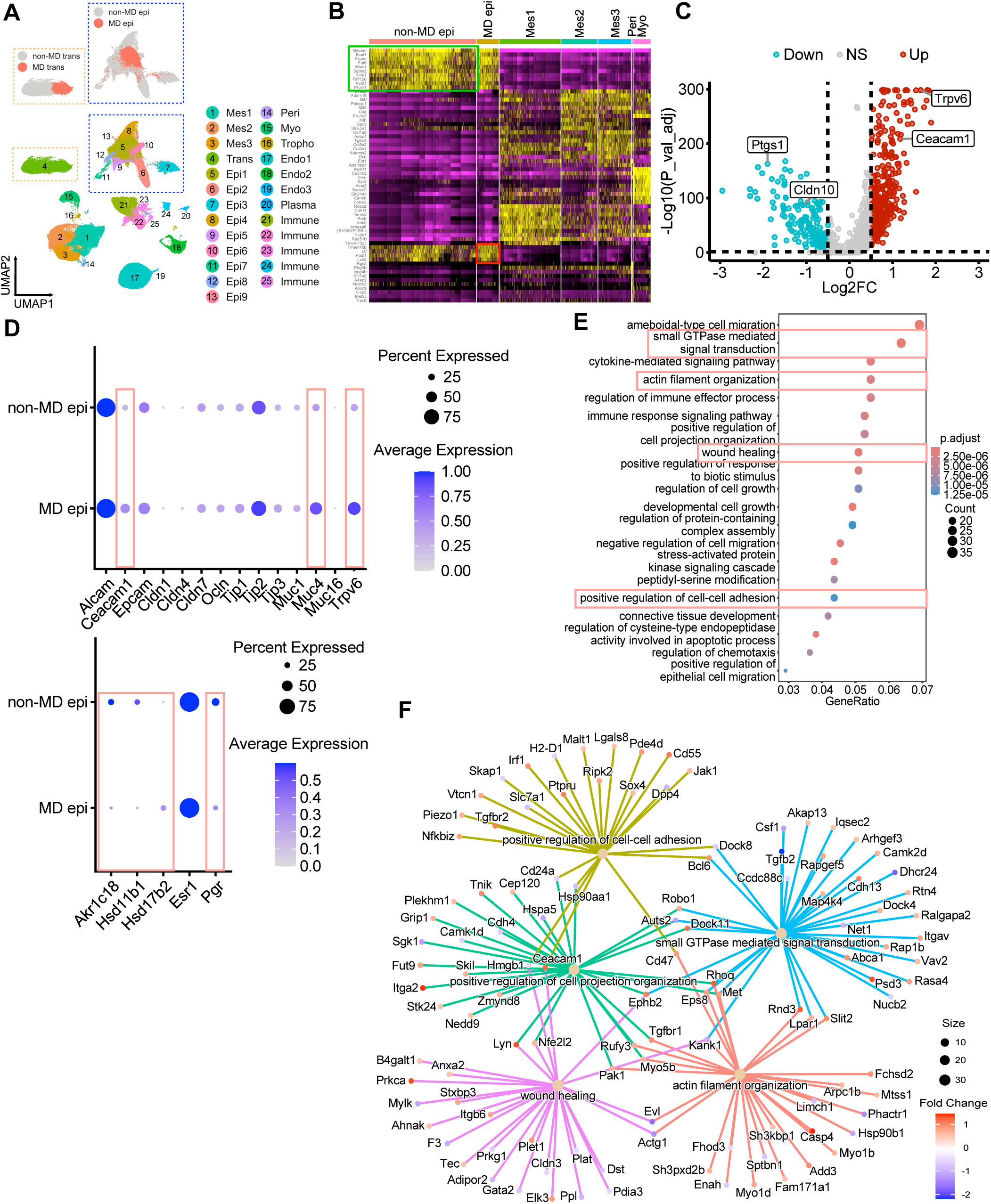
Transcriptional profiling indicates that mesenchymal-derived (MD) epithelial cells contribute to the barrier function of the epithelium. **(A)** UMAP of cluster annotation revealing 25 clusters, including 3 mesenchymal clusters (Mes1-3), 9 epithelial clusters (Epi1-9), 3 endothelial clusters (Endo1-3), 6 immune clusters (Immune1-5 and Plasma), 1 perivascular cluster (Peri), 1 myometrial cluster (Myo), and 1 transitional cluster (Trans) identified in the uteri of *Pdgfrα^CreERT2/+^; Rosa26-tdTomato^fl/+^* mice (composite of all time points E18.5 and PPDs1-28). The epithelial (blue dashed boxes) and transitional (orange dashed boxes) clusters were subset into non-MD epithelial/transitional and MD epithelial/transitional based on tdTom expression. **(B)** Heatmap of top 10 differentially expressed genes (DEGs) for non-MD epi (green box), MD epi (red box), Mes1, Mes2, Mes3, Peri, and Myo clusters. **(C)** Volcano plot showing 577 DEGs in MD epithelial cells compared to non-MD epithelial cells. **(D)** Dot plot of genes involved in cell-cell adhesion and epithelial barrier formation (top plot), and steroidogenesis and hormone receptors (bottom plot) in non-MD and MD epithelial cells. Red boxes indicate DEGs in MD epithelial cells. **(E)** Gene ontology analysis of enriched pathways in MD epithelial cells. Red boxes highlight pathways of interest. **(F)** Cnetplot showing representative pathways and associated genes in MD epithelial cells from (E) (red boxes).

Similarly, depletion of TRPV6, a membrane Ca^2+^ channel, was associated with impaired cell-cell contact formation (23), suggesting that *Trpv6* expression in MD epithelial cells may facilitate the acquisition of an epithelial phenotype by promoting cell-cell contact (**Fig. 2D**). *Muc4,* a cell surface mucin important for providing a protective barrier against infection, was also significantly up-regulated in MD epithelial cells (**Fig. 2D**). Gene ontology (GO) analysis of DEGs in MD epithelial cells showed enriched pathways associated with small GTPase-mediated signal transduction, actin filament organization, and positive regulation of cell-cell adhesion, all of which are involved in cellular morphology and transitions, and in wound healing, another enriched pathway (**Fig. 2E and F**). Notably, estrogen receptor alpha (*Esr1*) was abundantly expressed in MD and non-MD epithelial cells, but progesterone receptor (*Pgr*) was significantly decreased in MD epithelial cells (**Fig. 2D**). Furthermore, enzymes involved in steroid hormone synthesis and metabolism, including *Hsd11b1, Akr1c18, and Hsd11b2* were differentially expressed in MD epithelial cells (**Fig. 2D**). These data suggest that MD epithelial cells display enhanced epithelial barrier function but exhibit altered hormone production and responsiveness compared to resident non-MD epithelial cells.

### MD epithelial cells have limited growth potential

At E18.5, a small percentage of MD epithelial cells was observed, which could be a minor progenitor population that expanded during regeneration. This expansion, rather than *de novo* MET, could explain the increased number of MD epithelial cells observed at PPD1 and PPD2 (**Fig. 1B**). To investigate this, EpCAM^+^tdTom^+^ MD epithelial cells and EpCAM^+^tdTom^-^ non- MD epithelial cells were sorted and cultured for organoid formation potential. On day 7 of culture, non-MD epithelial cells formed round and cystic organoids (**Fig. 3A**). In comparison, MD epithelial cells failed to form typical organoids but rather formed small cell clusters that were fewer in number (**Fig. 3B, C**). On day 14 of culture, some MD epithelial cell clusters demised while non-MD epithelial organoids continued to grow. As cells were cultured in low concentration in single-cell suspension, the organoid-forming efficiency could be calculated as an indicator of stem/progenitor cell activity. The organoid-forming efficiency was lower in MD epithelial cells at 0.093% compared to non-MD epithelial cells (0.555%; **Fig 3B**). Limited organoid-forming potential might be a result of reduced cell proliferation. To evaluate this, we examined LE proliferation on PPD1 in uterine tissue sections and determined that the percentage of Ki67^+^ MD epithelial cells was significantly lower than Ki67^+^ non-MD epithelial cells (**Fig. 3D and E**). To corroborate this finding, we assessed a common proliferation marker gene set and found the majority of the genes were highly expressed in non-MD epithelial cells compared to MD epithelial cells (**Fig. 3F**). These data indicate that MD epithelial cells have limited growth potential and are likely more differentiated than non-MD epithelial cells.

**Figure 3.**
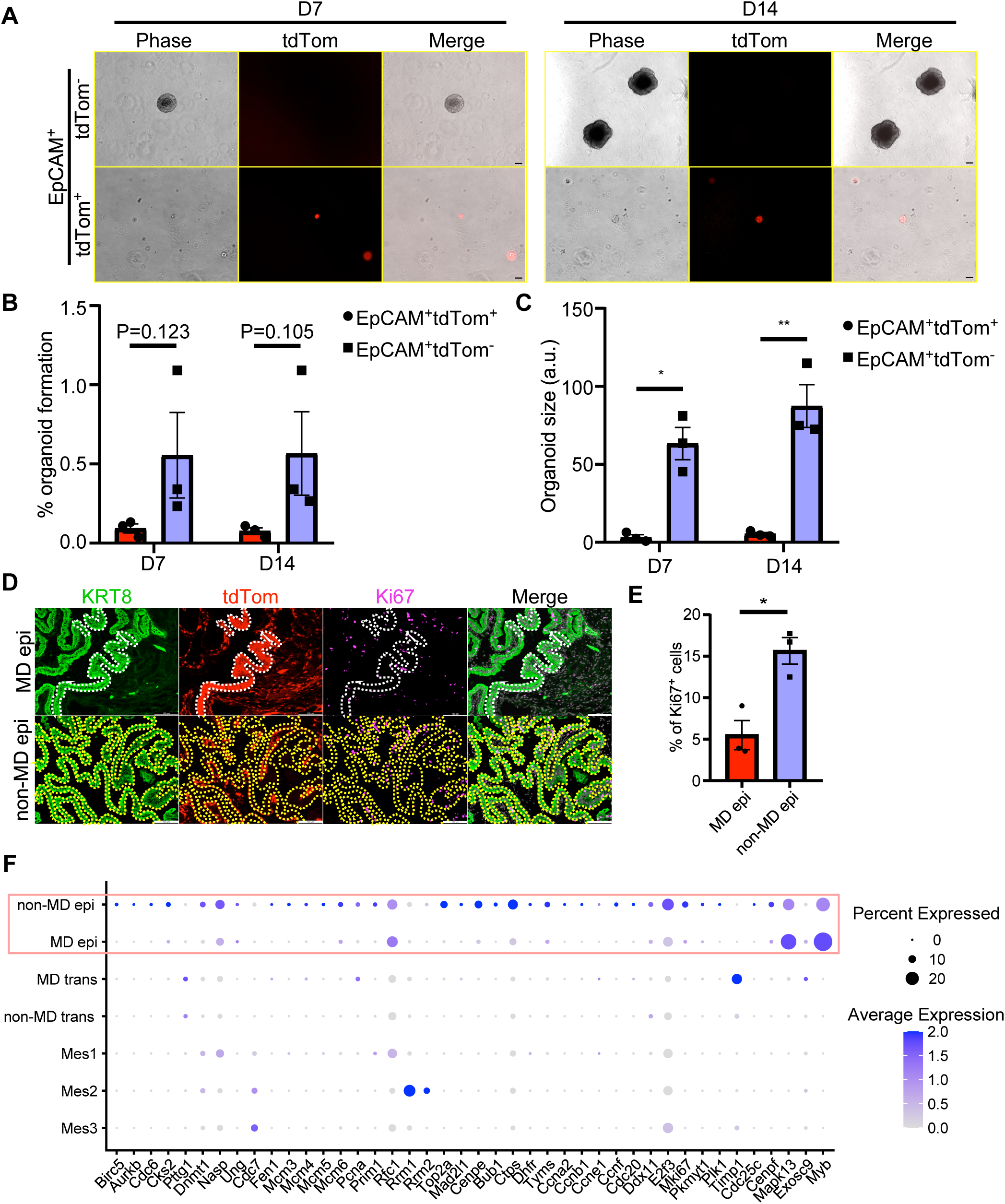
Mesenchymal-derived (MD) epithelial cells have reduced growth potential. **(A)** EpCAM^+^tdTom^-^ (non-MD, top panel) and EpCAM^+^tdTom^+^ (MD, bottom panel) endometrial epithelial cells were isolated and sorted from PPD1 *Pdgfrα^CreERT2/+^; Rosa26-tdTomato^fl/+^* mice uteri and cultured at 2000 cells/20 μl drop in Cultrex BME for 7 and 14 days. Scale bar: 100 µm. **(B)** Quantification of the number of organoids formed at day (D) 7 and D14. **(C)** Quantification of the size of organoids at D7 and D14 of culture. **(D)** Immunofluorescent staining for KRT8 (green) and Ki67 (magenta), with endogenous tdTom (red), and nuclear counterstaining with DAPI (grey, Merge) in MD (upper panel) and non-MD (lower panel) epithelial cells from uterine sections at PPD1. Dashed lines demarcate MD (yellow) and non-MD (white) epithelial cells for Ki67 expression for quantification. Scale bar: 100 µm. **(E)** Quantification of Ki67^+^ cells in MD and non-MD epithelial cells. **(F)** Dot plot of a series of proliferation genes in different cell types from snRNA-seq. Red box highlights non-MD and MD epithelial cells. * P < 0.05; ** P < 0.01.

### MD epithelial cells are transient and removed through mechanisms of regulated cell death (RCD)

We first looked into the cell composition at different time points during postpartum (24). The overall proportion of epithelial cells (MD and non-MD) decreased across postpartum regeneration (*SI Appendix,* **Fig. S3A**). This was due in part to apoptosis as evidenced by cleaved caspase 3 (CC3) immunofluorescence in tissue sections (**Fig. 4A**) and enrichment for apoptosis-related genes by snRNA-Seq (**Fig. 4B**). Uniquely, the MD epithelial cells were dynamically regulated during regeneration, increasing rapidly after parturition then declining quickly after PPD2, indicating a preferential loss compared to non-MD epithelial cells.

**Figure 4.**
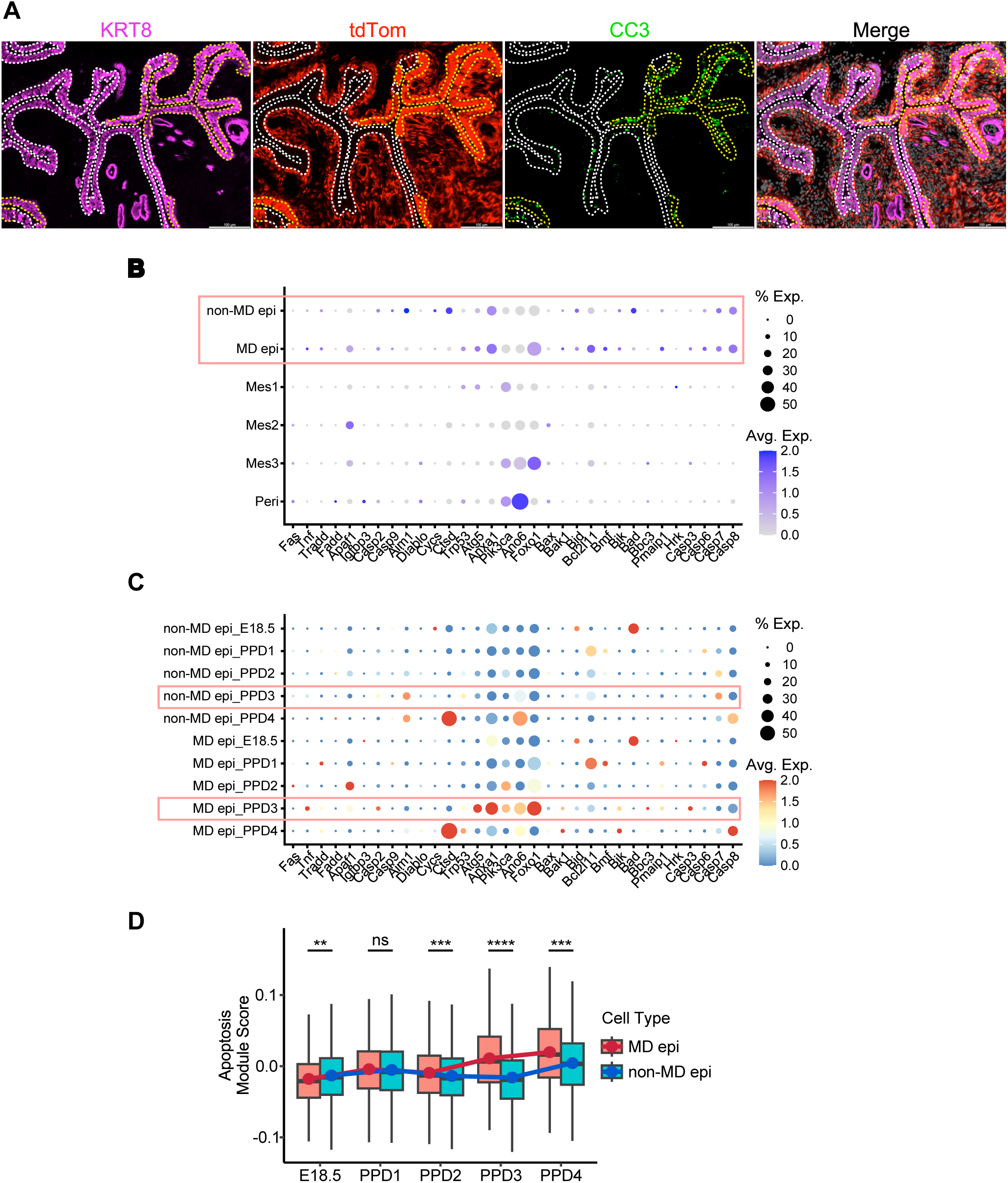
Mesenchymal-derived (MD) epithelial cells are removed via apoptosis. **(A)** Immunofluorescent staining for KRT8 (magenta) and cleaved caspase3 (CC3, green), with endogenous tdTom (red), and nuclear counterstaining with DAPI (grey, Merge) in *Pdgfrα^CreERT2/+^; Rosa26-tdTomato^fl/+^* mice uteri on PPD2. Non-MD epithelial cells are outlined by white dashed lines, and the MD epithelial cells are outlined by yellow dashed lines. **(B)** Dot plot of pro-apoptosis genes in different cell clusters. Red box highlights non-MD and MD epithelial cells. **(C)** Dot plot of pro-apoptosis genes in non-MD and MD epithelial cells at E18.5 and PPDs1-4. Red boxes highlight differential gene expression at PPD3. **(D)** Box plot of module scores for pro-apoptosis genes at specified time points during endometrial regeneration. A red and blue trend line connecting the mean of the module scores was created for MD and non-MD epithelial cells, respectively. ** P < 0.01, *** P < 0.001, **** P < 0.0001; ns, not statistically different.

Apoptosis-related genes were then evaluated in MD and non-MD epithelial cell clusters across regeneration time points. Notably, expression levels of a series of pro-apoptosis genes began increasing on PPD1 with peak expression of specific genes including *Tnf, Caps2, Casp3, Atg5, Anxa1, Pik3ca*, *Ano6* and *Foxo1* on PPD3 in MD epithelial cells but not in non-MD epithelial cells (**Fig. 4C and 4D**). We further investigated other RCD mechanisms (*SI Appendix* **Dataset S2**) including anoikis (a specialized type of apoptosis when cells lose attachment to the extracellular matrix and neighboring cells; *SI Appendix,* **Fig. S3B, C and J**), pyroptosis (gasdermin-mediated programmed death; *SI Appendix,* **Fig. S3D, E and K**), ferroptosis (cell death characterized by iron-dependent lipid peroxidation; *SI Appendix,* **Fig. S3F, G and L**), and autophagy (*SI Appendix,* **Fig. S3H, I and M**). Overall, we observed a similar pattern but elevated level of different RCDs in MD epithelial cells compared to non-MD epithelial cells, suggesting higher susceptibility of MD epithelial cells to programmed cell death, reinforcing the transient nature of MD epithelial cells.

Collectively, these data suggest that MET is triggered in response to wounding to help facilitate the rapid restoration of the endometrial epithelial barrier, followed by replacement of MD epithelial cells through various mechanisms of RCD.

### Transitional cells undergoing MET were identified in the regenerating endometrium

In our early studies using the menses-like mouse model (11) of endometrial regeneration, we characterized a population of pan-KRT and vimentin (mesenchymal marker) co-expressing “stromal transitional cells” in the mesometrial zone of regeneration. Similarly, in the current pregnancy model, we observed KRT8^+^ cells residing in the mesometrial stroma on PPD1 that appeared to migrate to the regenerating epithelium (**Fig. 5A, C**). To assess the KRT8^+^ stromal mesenchymal cells as potential “transitional” cells undergoing MET, we first characterized their cellular identity by immunofluorescence. Many of the KRT8^+^ stromal cells expressed RFP (*i.e.* tdTom) (*SI Appendix,* **Fig. S4A**) and were negative for CD45 (pan-immune cell marker; *SI Appendix,* **Fig. S4B**), indicating their mesenchymal phenotype. Notably, KRT8^+^ cells within the stromal compartment did not exhibit expression of EpCAM (*SI Appendix,* **Fig. S4C**) or CDH1 (**Fig. 5D**), both of which are critical for epithelial cell-cell adhesion. However, a rare subset of KRT8^+^ cells co-expressing CDH1 was observed at the LE, where they appeared to be integrating into the epithelial layer (**Fig. 5D**, inset). Quantification of these potential transitional cells (EpCAM^-^tdTom^+^KRT8^+^) in the undecidualized stroma (EpCAM^-^tdTom^+^) by flow cytometry showed the highest percentage on PPD1 (13.57%; **Fig. 5B**). We reasoned that if these cells represent transitional mesenchymal cells involved in re-epithelialization, they would emerge at parturition and undergo MET, ultimately differentiating into MD epithelial cells during postpartum re-epithelialization. Consequently, the number of transitional cells would be expected to peak prior to the peak of MD epithelial cells on PPD2, a pattern consistent with observations from flow cytometry analysis (**Fig. 5E**). Importantly, we observed a similar pattern from the snRNA-seq data (**Fig. 5F**). During MET, mesenchymal cells lose their migratory ability as they establish cell-cell adhesions and attach to the basement membrane (7). These data suggest that KRT8^+^ transitional cells located in the stroma migrate to the LE where they complete MET and incorporate into the LE, upregulating epithelial adhesion proteins.

**Figure 5.**
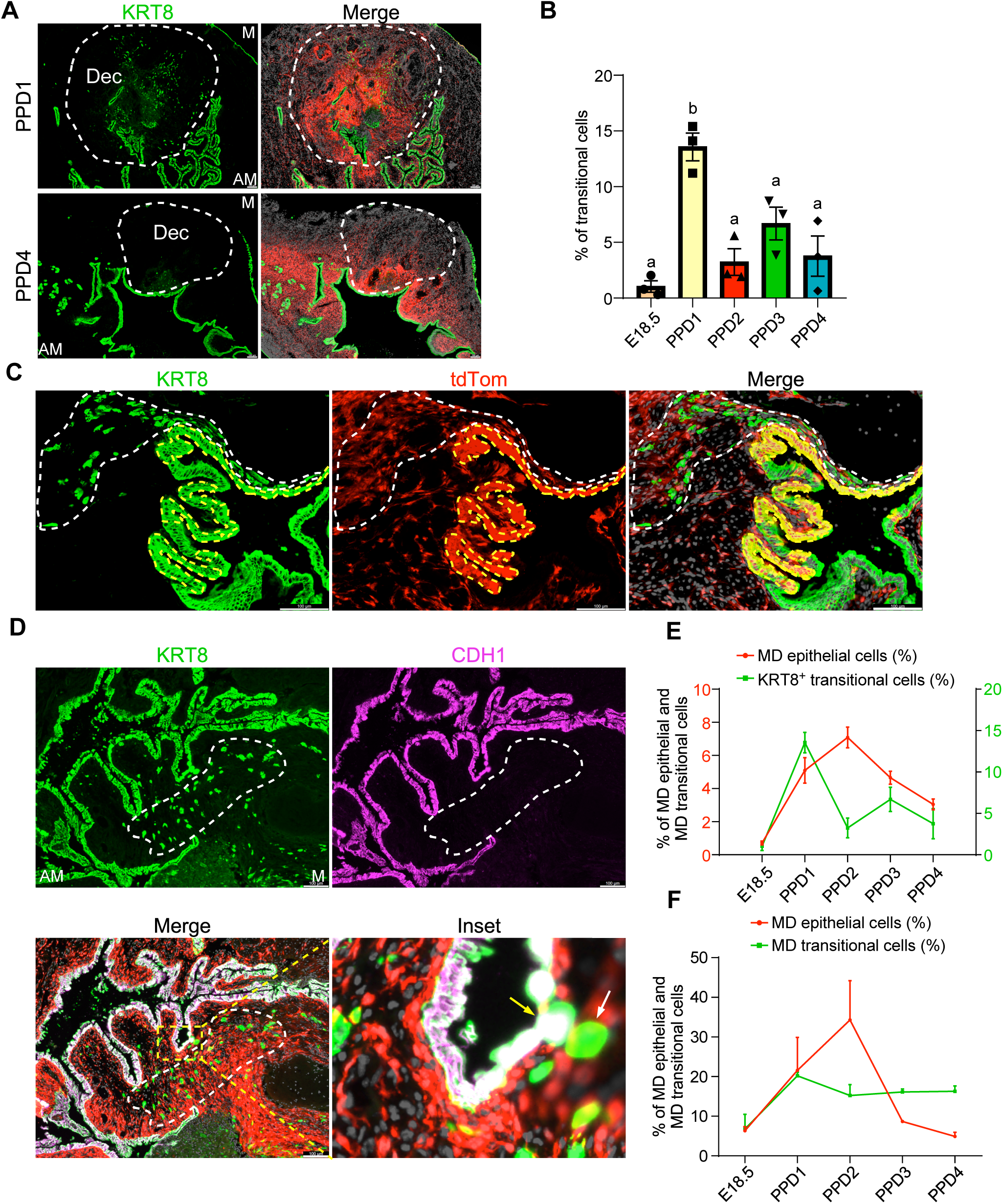
Mesenchymal-derived (MD) transitional cells were identified in the regenerating endometrium. **(A)** Immunofluorescent (IF) staining for KRT8 (green), endogenous tdTom (red), and nuclear counterstaining with DAPI (grey, Merge) in uteri of *Pdgfrα^CreERT2/+^; Rosa26-tdTomato^fl/+^*mice showing a population of KRT8^+^ transitional cells in the mesometrial (M) decidua (Dec, dashed white area) at PPD1 and PPD4. AM: anti-mesometrial pole; scale bar: 100 µm. **(B)** Quantification of the percentage of MD transitional cells in the endometrium of postpartum mice by flow cytometry analysis. MD-transitional cells were identified as EpCAM^-^ tdTom^+^KRT8^+^ and quantified as a percentage of the tdTom-labeled stromal cells (EpCAM^-^ tdTom^+^). **(C)** IF staining for KRT8 (green), endogenous tdTom (red), and nuclear counterstaining with DAPI (grey, Merge) showing migratory KRT8^+^ transitional cells (dashed white line) near MD epithelial cells (dashed yellow areas) in the regenerating endometrium on PPD1. **(D)** IF staining for KRT8 (green) and CDH1 (magenta), with endogenous tdTom (red) and nuclear counterstaining with DAPI (grey, Merge) in a PPD1 uterus. The white dashed area shows KRT8^+^CDH1^-^ transitional cells. Inset is a zoomed-in image of the yellow dashed box in the merged image. The yellow arrow indicates a cell co-expressing CDH1 and KRT8 and incorporating into the luminal epithelium. The white arrow indicates a cell expressing KRT8 but not CDH1. **(E)** Comparison of the dynamic change of MD epithelial cells and MD transitional cells by flow cytometry. % MD epithelial cells = EpCAM^+^tdom^+^/EpCAM^+^; % MD transitional cells = EpCAM^-^tdTom^+^KRT8^+^ /EpCAM^-^tdTom^+^. Note that the peak of transitional cells precedes the peak of MD epithelial cells. **(F)** Comparison of the dynamic change of MD epithelial cells and MD transitional cells by snRNA-seq. Percentages were calculated as *tdTom^+^* epithelial cells over total epithelial cells (Epi1-9) and *tdTom^+^* transitional cells over total non-epithelial *tdTom^+^* cells.

### snRNA-seq of postpartum regenerating endometrium confirms the presence of transitional cells undergoing MET

To gain insights into the molecular process of MET, we analyzed snRNA-seq data on uteri at all time points. As mentioned above, a distinct cluster (**Fig. 2A**, cluster 4) that expressed both mesenchymal and epithelial marker genes was identified and postulated to be the transitional cells observed by immunofluorescence and flow cytometry. Not all cells in the transitional cluster 4 expressed tdTom showing similar heterogeneity seen by immunofluorescence and flow cytometry. To refine this population, cells were subclustered by tdTom expression to identify those transitional cells that were mesenchyme derived (MD). The two subclusters were designated as MD and non-MD transitional cells. The MD transitional population expressed mesenchymal genes such as *Col1a1, Col3a1, Cdh11,* and *Vim*, and highly expressed epithelial marker genes such as *Krt8, Krt18, Cd9, Clu,* and *Sdc4* (**Fig. 6A)**.

**Figure 6.**
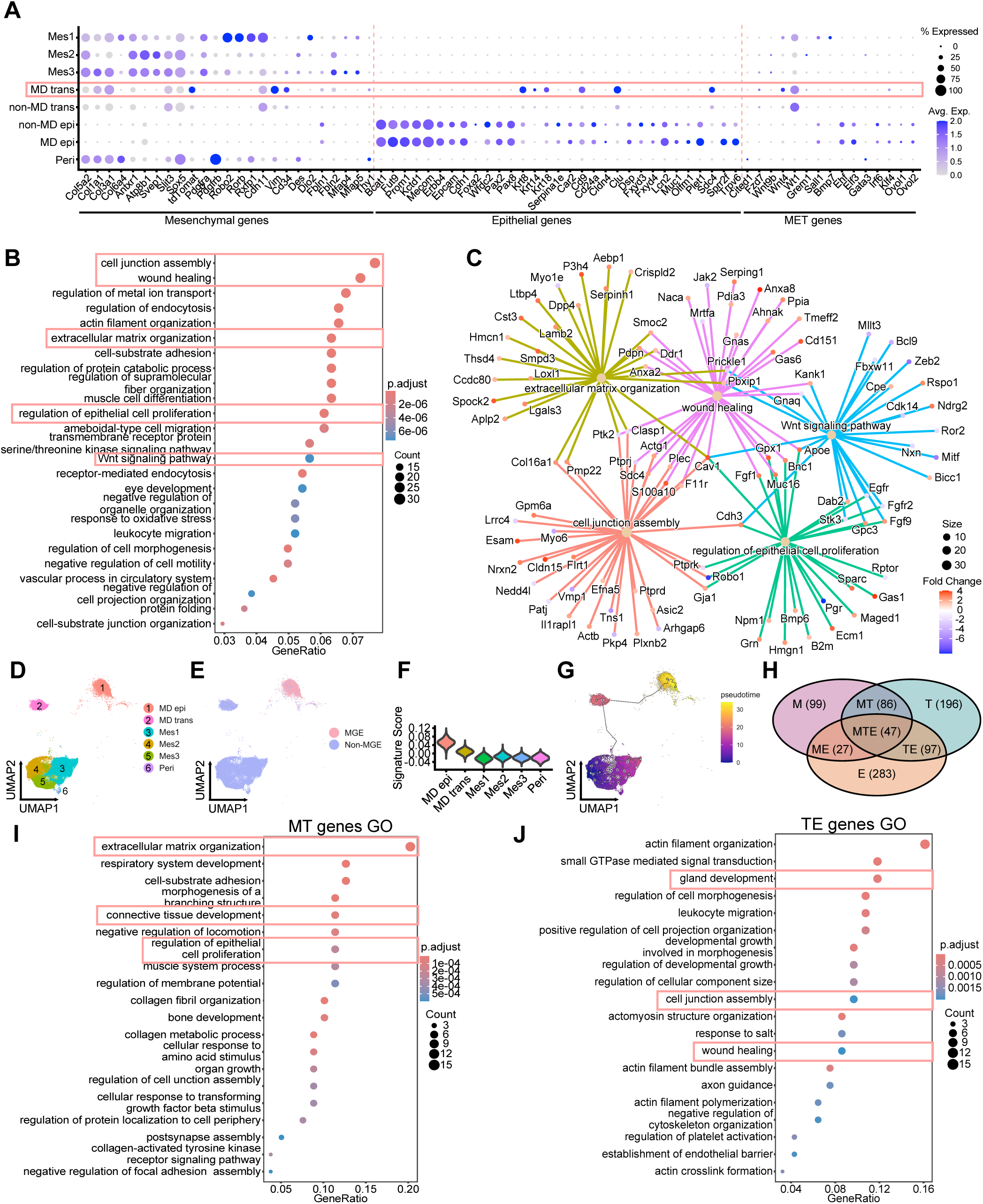
Characterization of mesenchymal-derived (MD) transitional cells by snRNA-seq. **(A)** Dot plot showing mesenchymal, epithelial and transitional cell cluster expression of mesenchymal, epithelial, and MET marker genes in the uteri of *Pdgfrα^CreERT2/+^; Rosa26-tdTomato^fl/+^* mice (aggregate of all time points). Red box highlights MD transitional cluster (MD trans). Red dashed lines demarcate gene sets. **(B)** Gene ontology (GO) analysis of enriched pathways in MD transitional cells. Red boxes highlight pathways of interest. **(C)** Cnetplot showing highlighted pathways from (B) and associated genes from GO analysis in MD transitional cells. **(D)** UMAP of mesenchymal cells (Mes1-3), MD transitional cells, MD epithelial cells, and perivascular cells. **(E)** AUCell analysis showing enrichment of a MET gene set in the MD epithelial cells. MGE: MET gene enriched. **(F)** MET gene set signature score in selected cell types. **(G)** Trajectory inference showing a root from mesenchymal cells through MD transitional cells to MD epithelial cells. **(H)** Venn diagram of the Monocle3 DEGs as a function of pseudotime. Numbers represent genes enriched in mesenchymal cells (M), MD transitional cells (T), MD epithelial cells (E) or shared amongst different cell types (MT, ME, TE and MTE). **(I)** GO of biological processes (BP) of enriched pathways in MT gene set. **(J)** GOBP of enriched pathways in TE gene set.

Notably, they were negative for *Epcam* and *Cdh1*, which was consistent with immunostaining from tissue sections, suggesting they were not mature epithelial cells and retained a migratory mesenchymal phenotype.

Studies have suggested that putative endometrial mesenchymal stem cells (eMSC) may undergo MET during endometrial epithelial regeneration (14). To clarify the stemness of MD transitional cells, we compared putative eMSC and endometrial epithelial stem cell (eESC) marker genes (a curated list from human and mouse studies) between perivascular, epithelial (MD and non-MD), MD transitional, and mesenchymal cells (*SI Appendix,* **Fig. S5A**). As expected, perivascular cells, proposed to contain eMSCs, expressed *Mcam*, *Pdgfrb, Itgb1, Thy1,* and *Cspg4* (*SI Appendix,* **Fig. S5A**). In comparison, MD transitional cells expressed *Itgb1, Aldh1a1*, *Cd34*, and *Klf4* (*SI Appendix,* **Fig. S5A**). Of note, CD34^+^KLF4^+^ putative eMSCs were shown to contribute to endometrial epithelial regeneration and repair (14). Putative eESC markers including *Axin2, Sox9*, and *Lgr5* were not expressed in the MD transitional population, however *Aldh1a1* was expressed. Since epithelial stem cells are more extensively studied in the small intestine, we referred to two reports characterizing a revival stem cell (revSC) population in the crypt (25) and an atrophy-induced villus epithelial cell (aVEC) population (26). We found the overall gene signatures for revSCs (38 genes) and aVECs (96 genes) were highly enriched (25.88-fold enrichment, P < 2.2e-16 for revSCs and 20.22-fold enrichment, P < 2.2e-16 for aVECs) in MD transitional cells compared to other cell types (*SI Appendix* **Fig. S5D and E, Dataset S3**), suggesting a progenitor epithelial cell state in MD transitional cells. To account for the apparent discrepancy to the endometrial stem cell analysis, we looked into the revSC and aVEC gene lists and found that these un-curated lists included genes that are associated with epithelial and cytoskeletal components, cell cycle and proliferation, and inflammatory and stress response, among others, suggesting that the MD transitional cells are responsive to wounding and inflammation and are actively undergoing cytoskeletal remodeling. Taken together, these data suggest that MD transitional cells are in a less-differentiated state allowing for transdifferentiation into epithelial cells.

To assess active MET in MD transitional cells, expression of key genes known to be involved in MET from the Molecular Signatures Database (MSigDB) (27) were examined.

Results show that MD transitional cells expressed several MET-associated genes, including *Wnt4, Wt1,* and *Klf4* (**Fig. 6A**). Gene ontology (GO) analysis of MD transitional cells revealed pathways that are related to tissue regeneration, including cell junction assembly, wound healing, extracellular matrix organization, and regulation of epithelial cell proliferation (**Fig. 6B and C**, *SI Appendix* **Dataset S4**). To further evaluate MET gene signatures, we referred to a study interrogating cholera toxin (Ctx)-induced MET in NAMEC8 human mammary epithelial-derived mesenchymal cells (28). Up-regulated genes were extracted (*SI Appendix* **Dataset S5**) and superimposed on our single-nuclei data set using AUCell to identify cells enriched for MET-associated genes. Both MD epithelial cells and MD transitional cells were enriched for MET genes above other cell types (**Fig. 6D-F**).

Using Monocle3 to perform trajectory inference analysis, we subset MD epithelial, MD transitional, mesenchymal 1-3, and perivascular cells (**Fig. 6D**) and found one partition suggesting a transdifferentiation may exist among different cell types. A starting root was programmatically determined in mesenchymal cell-2 cluster traveling through the MD transitional cell cluster and ending in the MD epithelial cell cluster (**Fig. 6G**). To further reveal the temporal transcriptional regulation of MET, we again took advantage of Monocle3 which allows for characterizing genes that change as a function of pseudotime. We identified 1025 genes that were scored as highly significant and categorized them into the following expression profiles: genes that were only enriched in mesenchymal cells (designated as M); genes that were only enriched in transitional cells (designated as T); and genes that were only enriched in MD epithelial cells (designated as E). Likewise, four more categories were assigned as MT (mesenchymal and MD transitional), TE (MD transitional and MD epithelial cells), ME (mesenchymal and MD epithelial cells), and MTE (mesenchymal, MD transitional and MD epithelial cells) (**Fig. 6H**, *SI Appendix* **Dataset S6**). MET was broadly segregated into two phases: phase one being mesenchymal (M) to transitional (T) state, and phase two being transitional (T) to epithelial (E) state. Accordingly, we investigated the MT and TE genes by gene set enrichment analysis. Enriched gene ontology of biological process (GOBP) terms for the MT gene set included extracellular matrix organization, connective tissue development, and regulation of epithelial cell proliferation (**Fig. 6I**), while enriched GOBP terms for the TE gene set included gland development, cell junction assembly, and wound healing (**Fig. 6J**). Together, snRNA-Seq of postpartum regenerating uteri confirmed the presence of MD transitional cells expressing mesenchymal and epithelial markers, which appear to be actively undergoing MET to contribute to postpartum endometrial regeneration.

### Cell-cell communication analysis predicts key signaling pathways involved in MET

The induction of MET is not well understood; therefore, we explored signaling pathways that may contribute to MET using CellChat (29). 50 signaling pathways were identified as biologically significant during the window of endometrial regeneration (E18.5 to PPD4) among MD and non-MD epithelial, MD transitional, mesenchymal, and perivascular cells. MD transitional cells were the most highly interactive population as they contributed most of the outgoing and incoming signals, while MD epithelial cells and perivascular cells were dominant signal receivers (**Fig. 7A**). Specifically, some of the prominent outgoing signaling networks of interest were EPHA, WNT, ncWNT and BMP (**Fig. 7B and C**). The canonical WNT pathway is crucial for embryonic development and adult tissue regeneration (30, 31). In our data, the most prominent interactions for WNT signaling were between the *Wnt4* ligand-expressing MD transitional cells and the *Fzd6/Lrp5/6* receptor-expressing MD epithelial cells (**Fig. 7D and E**). WNT4 is well known for its essential role in stromal cell decidualization, a type of MET. Compared to WNT signaling, ephrin-EPH signaling was more intricate involving more cell types, but the strongest interactions still remained between MD transitional cells and MD epithelial cells (**Fig. 7F and G**). More specifically, MD transitional cells expressed the ligand ephrin A5 (*Efna5*), while MD epithelial cells expressed the receptors *Ephb2* and *Epha4* (**Fig. 7G**). It has been reported that EPHA signaling plays a key role in somite morphogenesis by promoting mesenchymal cells in the paraxial mesoderm to differentiate into epithelial cells which delineate boundaries between somites (32, 33). Taken together, cell-cell communication analysis provides insights into potential key pathways and factors driving MET during postpartum endometrial regeneration.

**Figure 7.**
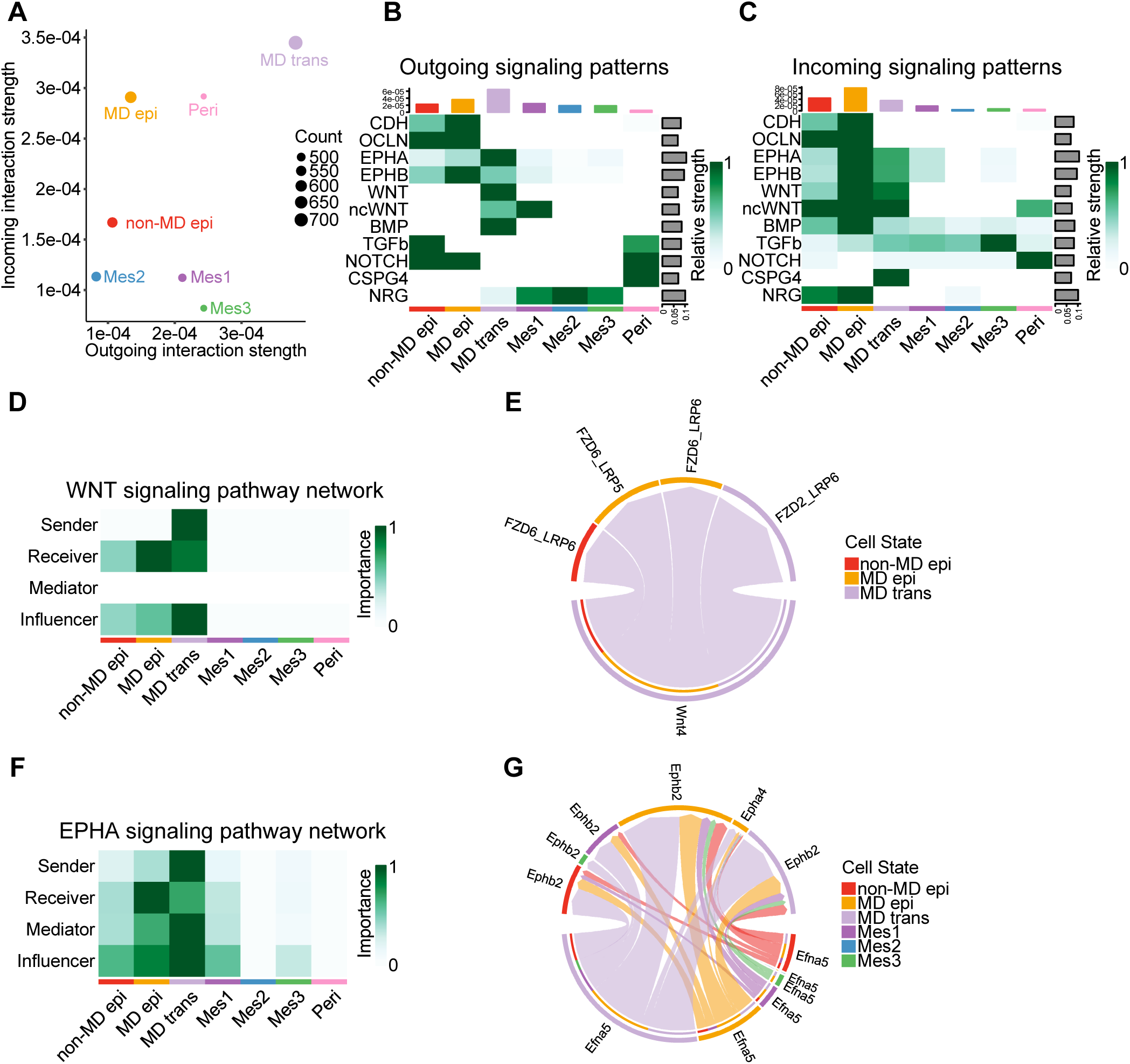
Cell-cell communication analysis reveals strong interactions between mesenchymal-derived (MD) transitional and epithelial cells. **(A)** Scatter plot showing aggregate outgoing and incoming signals among indicated cell clusters from *Pdgfrα^CreERT2/+^; Rosa26-tdTomato^fl/+^* mice (composite of all time points) **(B)** Heatmap showing outgoing signal patterns of interest. **(C)** Heatmap showing incoming signal patterns of interest. **(D)** Heatmap of the WNT signaling pathway showing the relative importance of each cell type based on the four network centrality measures. **(E)** Chord plot of the ligand-receptor interactions of the WNT pathway. The lower segment shows the cell type(s) expressing the ligands and the upper segment shows the cell type(s) expressing the receptors. The size of the outer bars indicates the strength of the signal. **(F)** Heatmap of the EPHA signaling pathway. **(G)** Chord plot of the EPHA signaling pathway.

## Discussion

MET was first proposed as a mechanism for endometrial re-epithelialization by Walter Heape in 1897 (17); however, experimental evidence supporting this hypothesis did not emerge until over a century later (11, 12). While subsequent studies have largely supported the role of MET in endometrial repair, the hypothesis has also faced scrutiny. An alternative explanation posited that mesoepithelial cells of the embryonic Müllerian duct epithelium—the precursor to adult endometrial epithelium—were the source of mesenchymal promoter-labeled epithelial cells in the adult, rather than MET (16). Our investigation of postpartum endometrial re-epithelialization, together with a prior study examining menses-like repair (15), employed an inducible lineage-tracing mouse model (*Pdgfrα^CreERT2/+^; Rosa26-tdTomato^fl/+^*) to avoid confounding by embryonic labeling. These findings collectively provide confirmation that MET occurs in the adult endometrium. In the menses-like mouse model, EpCAM⁺tdTom⁺ MD epithelial cells peaked at 48 hours and declined at 72 hours during endometrial regeneration (15). A similar temporal pattern was observed in the present study, with MD epithelial cells peaking at PPD2 and beginning to decline at PPD3. However, the menses-like model limits long-term assessment of MD epithelial cells due to ovariectomy-induced endometrial atrophy. To overcome this limitation, we employed the postpartum regeneration model with intact ovaries, allowing extended observation of endometrial repair. In this context, MD epithelial cells continued to decline at PPD4 and returned to baseline levels by PPD28. This decline occurred via apoptosis and potentially other forms of regulated cell death, supporting the transient nature of these cells. Notably, our findings suggest a novel role for MD epithelial cells in providing an initial epithelial barrier to rapidly cover the denuded endometrial surface, followed by their replacement with resident non-MD epithelial cells.

A key advantage of the current mouse model is the ability to identify MD epithelial cells via tdTom expression in snRNA-seq data, enabling direct comparison of their transcriptomic profiles to those of resident, non-MD epithelial cells. Through this approach, we found that MD epithelial cells contribute to the epithelial barrier function, characterized by expression of cell adhesion molecules (*e.g., Epcam, Alcam,* and *Ceacam*), cell junction proteins (*e.g., Tjps, Cldns*, *Ocln*, and *Cdh1*), and mucins (*e.g., Muc1* and *Muc4*). Notably, genes such as *Ceacam1* and *Muc4* were significantly up-regulated relative to non-MD epithelial cells. In contrast, MD epithelial cells displayed altered expression of genes involved in steroid metabolism (*Hsd11b1, Akr1c18, Hsd11b2*) and hormone signaling (*Pgr*), suggesting a divergence from the functional profile of typical endometrial epithelial cells. In addition, MD epithelial cells demonstrated reduced proliferative capacity *in vivo* and limited organoid-forming potential *in vitro*. Together with their transient presence during regeneration, these findings support the conclusion that MD epithelial cells primarily serve a temporary barrier function rather than participating in long-term epithelial maintenance.

Additional evidence supporting MET during endometrial re-epithelialization was obtained through the identification of transitional cells both *in vivo* and in the snRNA-seq dataset. Using tdTom expression to specifically trace the Pdgfrα-lineage, we identified a population of transitional cells and demonstrated a trajectory from mesenchymal cells through MD transitional cells to fully differentiated MD epithelial cells. Putative stromal stem/progenitor cells expressing CD34 and KLF4 (14) or Nestin (34) were proposed as the origin of mesenchymal cells undergoing MET. While we did not observe a trajectory originating from the perivascular cluster—previously identified as a candidate eMSC population (35)—MD transitional cells were enriched for *Cd34*, and to a lesser extent *Klf4*, suggesting they may represent a less differentiated state along this lineage. These findings raise the possibility that a subset of stromal progenitors contributes to the transitional cell pool during endometrial repair. A comparable transitional cluster was previously observed in the menses-like regeneration model using *Pdgfrb-BAC-eGFP* mice (15). Interestingly, transitional cells in the menses-like model expressed canonical epithelial markers such as *Cdh1* and *Epcam*. In contrast, transitional cells in our postpartum model lacked expression of CDH1 and EPCAM at both the RNA and protein levels, consistent with the migratory phenotype observed *in vivo*. These differences likely reflect the enhanced specificity afforded by tdTom expression, which enabled more precise delineation of the transitional cell population.

Further characterization of the transitional cell cluster revealed that *Wilms tumor 1* (*Wt1*) was highly expressed in MD (and non-MD) transitional cells. *Wt1* has been implicated in the regulation of MET and EMT in a tissue-specific context. For example, during kidney development, metanephric mesenchymal cells undergo MET to form epithelialized structures and ultimately give rise to mature nephrons (36). In this setting, *Wt1* is highly expressed and regulates key genes such as *Wnt4* and *Cdh1*, which drive epithelial differentiation. Loss or mutation of *Wt1* disrupts this process, resulting in impaired MET and developmental abnormalities such as Wilms tumor, characterized by the persistence of undifferentiated, non-epithelialized cells (36). In our dataset, *Wnt4*—which also plays a critical role in stromal cell decidualization, another MET-associated process—was abundantly and specifically expressed in MD transitional cells, further supporting their involvement in MET during endometrial regeneration. GOBP analysis of DEGs in MD transitional cells revealed significant enrichment of pathways involved in junction assembly, adhesion organization, and regulation of cell migration—processes consistent with a role in MET. Importantly, components of the WNT signaling pathway were also significantly overrepresented, further implicating this pathway in the regulation of transitional cell states during endometrial repair.

The mechanisms underlying the induction of MET in the endometrium remain poorly defined. To address this, we used CellChat analysis to investigate signaling pathways potentially contributing to MET during regeneration. Consistent with findings from GOBP analysis, WNT signaling emerged as a highly interactive pathway between MD transitional cells expressing *Wnt4* and MD epithelial cells, which expressed the receptors *Fzd6* and *Lrp5/6*. Notably, WNT4 signaling is essential for stromal cell decidualization, a process also characterized as a form of MET and is required for successful embryo implantation. In mice, decidualization is dependent on the WNT4–FZD–CTNNB1–FOXO1 axis (37), and disruption of *Wnt4* impairs this process (37, 38), suggesting a potential mechanistic continuity between MET events that occur during stromal cell decidualization and those involved in stromal cell-derived re-epithelialization.

Complementary evidence from human endometrial models supports this link: scRNA-seq of endometrial stromal and epithelial assembloids undergoing *in vitro* decidualization identified a transitional population enriched for GO terms associated with wound healing (39). A similar transitional cell population was observed in endometrial biopsies during the mid-to-late luteal phase, coinciding with decidualization *in vivo* (40). These findings have led to the intriguing hypothesis that transitional cells arising during decidualization may also contribute to endometrial re-epithelialization (41).

In addition to WNT signaling, our analysis suggests that ephrin signaling may represent a novel pathway promoting MET in endometrial re-epithelialization. Specifically, we identified *Efna5* expression in MD transitional cells, with corresponding *Epha4* and *Ephb2* receptors expressed in MD epithelial cells. EPHA signaling is known to regulate somite morphogenesis by promoting the MET of mesenchymal cells within the paraxial mesoderm. Activation of EPHA4 in this context induces apical redistribution of β-catenin (*Ctnnb1*), polarization, and acquisition of columnar morphology—hallmarks of epithelial differentiation (32, 33). Furthermore, EPHB2 was shown to suppress colorectal cancer progression, consistent with a role in antagonizing EMT (42). Of interest, *Epha4* has been detected in menstrual fluid (43), which contains a rich milieu of factors implicated in endometrial repair (44), and its expression is elevated in endometriotic lesions (45), suggesting a potential role for EPHA signaling in pathological endometrial remodeling. Together, these findings support ephrin signaling as a candidate regulator of MET during endometrial regeneration.

In summary, the current study presents several key findings: (1) MET is further substantiated as a mechanism contributing to endometrial re-epithelialization; (2) MD epithelial cells represent a distinct, transient population that facilitates rapid restoration of the epithelium; and (3) multiple candidate signaling pathways promoting MET were identified (**Fig. 8**). To our knowledge, this is the first report to characterize the unique transcriptomic signature of MD epithelial cells and delineate their differential regulation compared to resident, non-MD epithelial cells during endometrial re-epithelializatoin. Moreover, by leveraging tdTom expression to precisely define cell populations, we were able to directly trace the progression from mesenchymal to transitional to MD epithelial cells and identify associated signaling interactions that may drive MET.

**Figure 8.**
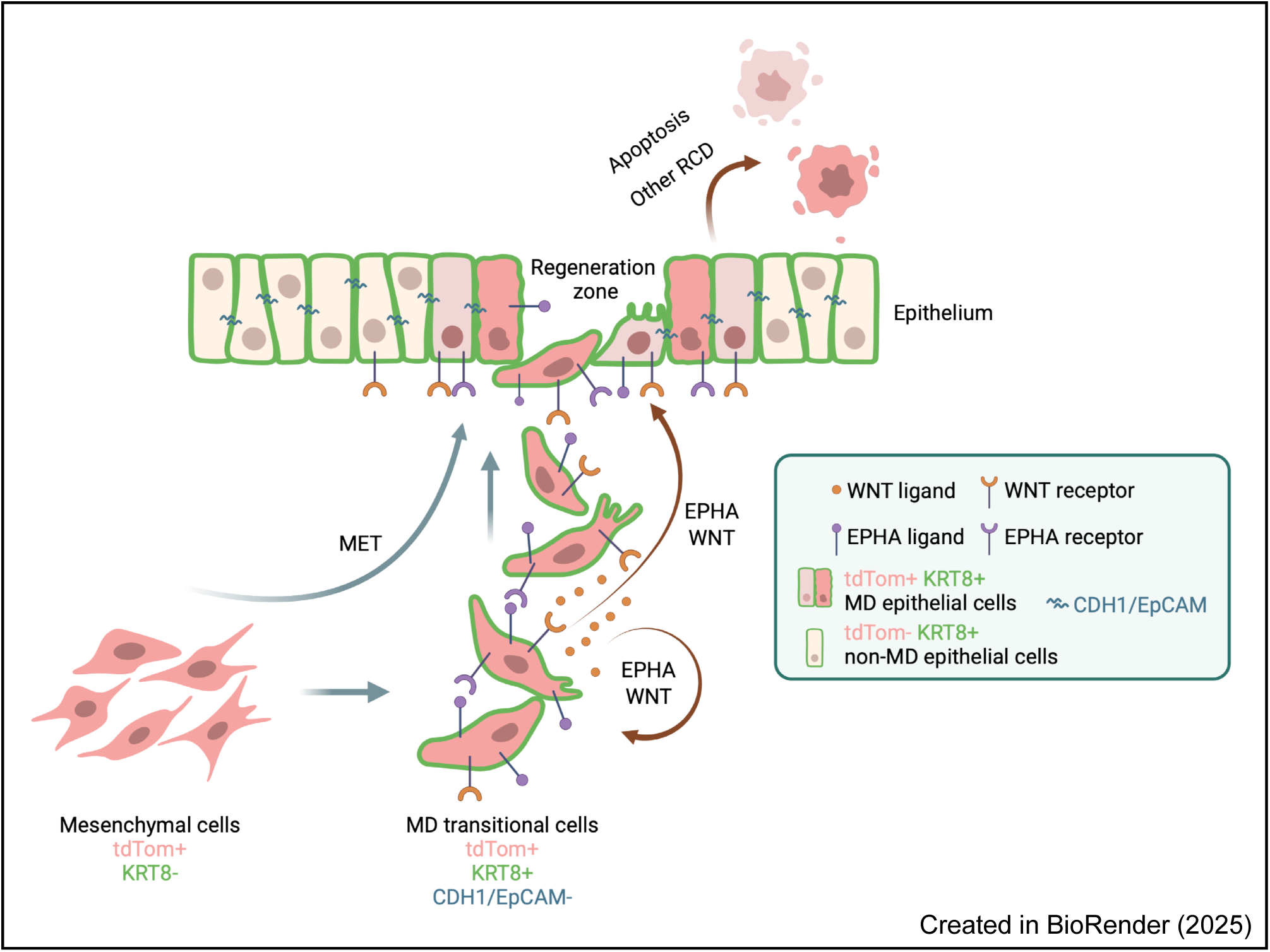
Proposed mechanism of MET during postpartum endometrial re-epithelialization. tdTom+ mesenchymal cells undergo partial MET to differentiate into KRT8+ MD transitional cells. MD transitional cells bear a dual phenotype of mesenchymal and epithelial features, migrating to the regeneration zone, completing MET to incorporate to the denuded luminal epithelium. E-cadherin (CDH1) and epithelial cell adhesion molecule (EpCAM) are induced upon incorporation into the epithelium. Potential key mediators of MET include WNT and EPHA signaling via autocrine and paracrine mechanisms. After timely restoration of the LE layer, the MD epithelial cells are subjected to various regulated cell death (RCD) mechanisms, mainly apoptosis, to be cleared and replaced by the native non-MD epithelial cells.

These findings have important implications for understanding human endometrial biology and disease. Dysregulation of MET may contribute to pathological conditions such as endometriosis, where misplaced endometrial cells exhibit aberrant plasticity, or endometrial cancer, in which epithelial-mesenchymal dynamics are often co-opted to promote tumor progression. The identification of specific signaling pathways involved in physiological MET may inform therapeutic strategies aimed at modulating cellular plasticity in these disease contexts.

Future studies are warranted to determine whether similar cellular trajectories and molecular signatures are present in human endometrium, particularly in the context of menstruation, postpartum repair, and disease states. Additionally, targeting MET-related pathways may represent a novel approach to enhance regenerative therapies or disrupt pathological remodeling in endometrial disorders.

## Materials and Methods

### Animals and lineage tracing model

All methods used in this study were approved by the University of Missouri Institutional Animal Care and Use Committee. *Pdgfrα^CreERT2/+^* mice (46) and *Rosa26-tdTomato^fl/fl^* reporter mice (47) were purchased from The Jackson Laboratory (stock # 032770 and # 007914, respectively). *Pdgfrα^CreERT2/+^* mice express a tamoxifen-inducible Cre recombinase-mutant estrogen receptor (ER) fusion protein under the control of the mesenchyme-specific *Pdgfrα* promoter. The resultant CreERT2 fusion protein will translocate to the nucleus only upon tamoxifen (TAM) treatment. *Rosa26-tdTomato^fl/fl^* mice contain a loxP-floxed stop sequence preventing downstream expression of tdTomato (tdTom). After Cre-mediated recombination, tdTom will be constitutively expressed in Pdgfrα-expressing mesenchymal cells and any cells derived therefrom.

*Pdgfrα^CreERT2/+^* and *Rosa26-tdTomato^fl/fl^* mice were crossed to generate *Pdgfrα^CreERT2/+^; Rosa26-tdTomato^fl/+^* lineage tracing females and littermate controls (*Pdgfrα^+/+^; Rosa26-tdTomato^fl/+^*; **Fig. S1A**). To induce tdTom expression, TAM (Sigma-Aldrich, St. Louis, MO; 2mg/20g body weight in sesame oil) was administered via intraperitoneal injection for 3 consecutive days, followed by a wash-out period of at least 28 days before entering an experiment. TAM is an estrogenic compound having adverse effects on the endometrium, including endometrial hyperplasia (48). To minimize such effects, we adopted a protocol from a previous study showing a 4-week washout period post-TAM treatment to have no deleterious effects on the mouse uterus (15).

### Tissue collection during the estrous cycle and postpartum repair

For tissue collection during the estrous cycle, *Pdgfrα^CreERT2/+^; Rosa26-tdTomato^fl/+^* and control females at the first estrus and diestrus stage after TAM washout were determined by vaginal lavage method using 10% trypan blue/PBS. The stage was determined by microscopic examination of the proportions of nucleated and cornified epithelial cells and leukocytes present.

Mice were sacrificed and uteri dissected. One piece of uterine horn was fixed for cryosectioning and the remainder of the uterus was processed for flow cytometry.

For tissue collection during postpartum repair *Pdgfrα^CreERT2/+^; Rosa26-tdTomato^fl/+^* and control females were mated with WT males and checked for the presence of a copulation plug each morning, which was designated day 0.5 of pregnancy. Uteri were collected at embryonic day (E)18.5, postpartum days (PPD) 1, 2, 3, 4, and 28. A segment of uterine horn with one fetus (E18.5, fetus removed, placenta intact) or previous implantation site (PPDs) was excised, fixed in 4% paraformaldehyde for 1h on ice, incubated in 15% sucrose overnight at 4°C and then embedded in Tissue-Tek OCT compound (Sakura Finetek) for cryosectioning and immunofluorescent staining. The remaining uterus was cut open along the broad ligament and fetuses and placentae (E18.5) or decidua (PPDs) were removed. A second uterine segment consisting of a single implantation site was snap frozen in liquid nitrogen for single-nuclei RNA sequencing (snRNA-seq) and the remainder of each uterus was further processed for flow cytometry analysis.

### Immunofluorescence

Frozen tissue sections were cut (8 μm) and mounted onto Superfrost Plus microscope slides (Fisher Scientific, Pittsburgh, PA) and dried on a heating station at 37°C for 3hrs. OCT was dissolved by incubating slides in PBS at 37°C for ∼10 min. Tissues were then blocked with PBS containing 0.1% Triton-X 100, 1% bovine serum albumin (BSA), and 5% normal goat serum for 1h at room temperature (RT), followed by overnight incubation at 4°C with primary antibodies (**Table 1**) diluted in PBS blocking solution. The following day, tissue sections were washed with PBS and incubated with secondary antibodies Alexa Fluor-488 goat anti-rabbit or Alex Fluor-647 goat anti-rat (1:1000; Molecular Probes, Eugene, OR) for 45 min at RT. After PBS washes, sections were counterstained with 300 nM DAPI (Life Technologies, Eugene, OR), and cover slipped using fluoro-gel aqueous mounting medium (Electron Microscopy Sciences, Hetfield, PA).

**Table 1.** Antibodies used in the study.

| Antibody name | Source | Catalog # | Species | Dilution |
| --- | --- | --- | --- | --- |
| Pdgfra | Cell Signaling | 3174 | Rabbit | 1:500 |
| Pdgfra | Invitrogen | 14-1401-82 | Rat | 1:100 |
| E-cadherin | Cell Signaling | 3195 | Rabbit | 1:800 |
| Cytokeratin 8 | DSHB | TROMA-I | Rat | 1:50 |
| EpCAM | Abcam | Ab71916 | Rabbit | 1:400 |
| Vimentin | Cell Signaling | 5741 | Rabbit | 1:100 |
| Ki-67 | Cell Signaling | 9129 | Rabbit | 1:400 |
| Cleaved caspase 3 | Cell Signaling | 9664 | Rabbit | 1:400 |
| RFP | Rockland<br>Immunochemicals | 600-401-379 | Rabbit | 1:500 |
| CD45 | Cell Signaling | 70257 | Rabbit | 1:100 |
| EpCAM (Alexa<br>Fluor® 488) | Biolegend | 118210 | Rat | 10 µl/10<sup>6</sup> cells |
| Cytokeratin 8 (Alexa<br>Fluor® 405) | Novas Biologicals | NB120-9287AF405 | Mouse | 3 µl/10<sup>6</sup> cells |

### Endometrial Cell isolation and flow cytometry

Endometrial epithelial and stromal cell isolation was performed using a modified protocol (10). Briefly, uteri cleared of fetuses/placentae (E18.5) or decidua (PPDs) were digested in 0.25% Trypsin in HBSS+ (HBSS plus 1x anti-biotic/mycotic) under the following conditions: 4°C with oscillation for 60 min, followed by RT for 40 min, then 37°C for 25 min. An equal amount of HBSS+ solution with 10% FBS was added to halt digestion. Cell suspension and tissue fragments were filtered through a 100 μm cell strainer and flowthrough was centrifuged at 500 g for 5 min. After centrifugation, red blood cells were lysed using ACK lysis buffer (Gibco, Grand Island, NY), followed by another wash with HBSS+ and centrifugation. The resulting epithelial/stromal cells were resuspended in PBS sorting buffer (2% FBS, 1% BSA, and 1mM EDTA) containing antibodies of interest (**Table 1**) at RT for 15 min protected from light. Following a wash with PBS sorting buffer, cells were stained with VivaFix Cell Viability dye (BioRad, Cat. #1351112) for 20 min at RT protected from light. After a final wash and resuspension with PBS sorting buffer, cells were sorted on a S3e Cell Sorter (405/488/561 nm lasers, BioRad, Hercules, CA) using Prosort (v1.6.0.12) software. Briefly, events were gated by forward and side scatter, for singlets (forward scatter height × area), and then for live cells (VivaFix-410^-^). Epithelial cells were gated as EpCAM^+^ then as tdTom^-^ (non-MD) or tdTom^+^ (MD), mesenchymal cells were gated as EpCAM^-^ and tdTom^+^, and transitional cells were gated as EpCAM^-^ tdTom^+^ then KRT8^+^ (*SI Appendix,* **Fig. S1C**). Unstained cells and fluorescence minus one (FMO) were used as gating controls. Data were analyzed with FlowJo software (v10.10.0. FlowJo, Ashland, OR).

### Organoid formation assay

MD (EpCAM^+^tdTom^+^) and non-MD (EpCAM^+^tdTom^-^) epithelial cells from PPD 1 uteri (n=3) were isolated, flow cytometry sorted and seeded at 2000 cells/20 μl drop in 80% basement membrane extract (R&D, Cat. #3433-010-R1). Organoids were cultured in mouse organoid expansion media (49) supplemented with 10 μM Y27632 for 14 days. The number of organoids were counted in all drops (n=3) on days 7 and day 14 and organoid formation was calculated as (# organoids formed / # cells seeded) * 100. The size of each organoid was measured on days 7 and 14 using ImageJ/Fiji (ImageJ 1.54g) (50).

### Quantification of proliferating cells

To quantify the percentage of Ki67^+^ cells in MD and non-MD epithelial cells, four random areas of uterine sections that contained both epithelial types were selected and imaged from three individual mice on PPD1. The total number of MD and non-MD epithelial cells in each image was calculated by counting the number of DAPI positive areas using Analyze Particles function in ImageJ, with particle size adjusted to 0.003-0.01 and circularity adjusted to 0.5-1.0. The number of Ki67^+^ MD and non-MD epithelial cells were counted by eye in each image and subsequently the percentage of Ki67^+^ MD or non-MD epithelial cells was calculated.

### snRNA-seq sample preparation

Nuclei were isolated from snap-frozen uterine tissues using the Singulator 100 System (S2 Genomics) according to the manufacturer’s instructions. Briefly, tissues were cut into 50-80 mg pieces, placed in the NIC+ (nuclei isolation cartridge; S2 Genomics, cat. # 100-063-732) with 10 μl of 40 U/μl RNase inhibitor (Sigma-Aldrich, St. Louis, MO). After running the standard isolation protocol, nuclei were centrifuged at 500 g for 5 min at 4°C, washed with nuclei storage buffer (NSR, plus 0.4 U/μl RNase inhibitor), centrifuged again, resuspended in 250 μl NSR, and passed through a 40 μm cell strainer. After imaging on a Cellometer (Nexcelom Bioscience, Lawrence, MA) to determine concentration and quality of the nuclei preparations, the samples were sent to the University of Missouri Genomics Technology Core (MUGTC). RNA library was constructed using Chromium Next GEM Single Cell 3ʹ Reagent Kits v3.1 (Dual Index) and sequencing was performed on the Illumina NovaSeq S4 -PE100 platform with a target count of 10,000 nuclei per sample and a sequencing depth of 50,000 read pairs per nucleus. Two individual mice were included for sequencing per time point.

### Bioinformatic analyses

The raw sequence data are deposited at GEO under accession # GSE296238. The untrimmed sequencing files were aligned to a customized mm10 reference genome using Cell Ranger (version 7.0.1) by 10x Genomics. Specifically, the coding sequencing of tdTomato and the WPRE sequence were added to the reference genome to enable identification of reads mapping to the tdTomato gene. Following alignment, the raw count matrices were input to CellBender (version 0.3.0.) (51) for ambient RNA decontamination. The filtered count matrices were loaded into Seurat (version 5.1.0.) (52) for further processing. First, the doublets/multiplets (∼10% of the total reads) were filtered out using scDblFinder (version 1.16.0) (53). Then, cells with less than 200 or more than 4000 unique feature counts were filtered. In addition, cells with more than 5% of mitochondrial genes and 7% of ribosomal genes were filtered. The thresholds were set for filtering mitochondrial and ribosomal genes based on the distribution of genes in the dataset, so that cells with excessive expression of these genes were excluded (*SI Appendix,* **Fig. S2**). Lastly, genes expressed in less than three cells were removed. After pre-processing, the UMI count matrices were normalized using sctransform (an optimized method to replace NormalizeData, ScaleData, and FindVariableFeatures). The datasets were then subjected to integration with Harmony (54) using the IntegrateLayers function. The integration procedure returns a single dimensional reduction that captures the shared sources of variance across multiple conditions and can be used for unsupervised clustering analysis. Here, the ElbowPlot function was used to determine the principal components and the FindNeighbors and FindClusters functions were used to cluster cells (resolution = 0.5, dims = 1:30). Differentially expressed genes (DEGs) in all clusters were calculated with the FindAllMarkers function (logfc.threshold = 1, min.pct = 0.05) and cluster annotation was carried out with canonical marker genes for a given cell type. Both positive and negative values of fold changes were calculated for DEG analysis.

Gene ontology (GO) analysis was performed on marker genes of tdTom^+^ MD epithelial cells (*SI Appendix* **Dataset S1**) in the epithelial clusters and MD transitional cells (*SI Appendix* **Dataset S4**) in the transitional cluster to characterize enriched pathways from the gene sets in the Mouse Molecular Signature Database (MSigDB) under the M5: GO subcollection focusing on biological process (BP).

To curate a MET gene set, we assessed the RNA-seq data in table S8 from Pattabiraman et. al. (28). We extracted the most upregulated genes after treatment of mesenchymal derivatives of immortalized human mammary epithelial cells (N8 cells) by filtering genes with log2FC >2 followed by a ranking based on the P value. The top 10% of genes were extracted yielding a total of 339 genes (*SI Appendix* **Dataset S5**) and the list of genes was superimposed to our dataset for identification of clusters associated with MET. We conducted literature review to curate gene sets for the eESC and eMSC stem cell markers (44, 55–57), proliferation markers (58), pro-apoptosis genes (59, 60), pro-anoikis genes (61, 62), pro-pryoptosis (63), pro-ferroptosis (64) and pro-autophagy genes (65) (*SI Appendix* **Dataset S2**). The revSC gene set was obtained from supplementary table S2 (cluster 18) from Ayyaz, et al. (66) after removing ribosomal genes (*Rpl* and *Rps*). The aVEC gene set was obtained from supplementary table S1 from Ohara, et al. (67). Specifically, aVEC-specific genes, YAP-dependent genes in aVECs, and aVEC-specific & YAP-dependent genes were extracted (*SI Appendix* **Dataset S3**). AUCell (68) was used to identify various gene set enrichment in our dataset. AUCell calculates the enrichment of a gene set as an area under the curve (AUC) across the ranking of all genes (based on their expression level) in a cell. Cells whose signature genes were disproportionately represented among their highest-expressed genes received high AUC scores and were assigned as MGE (MET gene-enriched); others that fell below the threshold were labeled Non-MGE. For the MET (339), revSC (38), aVEC (96), eESC (20), eMSC (16), pro-apoptosis (32), pro-anoikis (19), pro-pryoptosis (17), pro-ferroptosis (35), and pro-autophagy (37) signature genes, a gene signature score per cell was calculated using the AddModuleScore function in R.

Trajectory analysis was performed using Monocle3 (69). Briefly, the datasets were subset into mesenchymal cells, perivascular cells, MD transitional cells, and MD epithelial cells. After data preprocessing, a root was computationally determined using the *get_earliest_principal_node* function from all selected cells. Then the *order_cells* function was used to place cells based on their transcription profiles. Lastly, a trajectory graph was created based on pseudotime inference. The graph_test function was used to characterize genes that were differentially expressed as a function of pseudotime. The initial gene list was then filtered by status == OK, q_value ≤ 0.05, and absolute morans_I ≥ 0.2 to return a final list containing 1025 genes for downstream analysis. To test the Monocle3 DEGs for their expression in respective clusters, the expression level of each gene was calculated in mesenchymal cells (M), MD transitional cells (T) and MD epithelial cells (E). Genes exceeding the threshold expression (threshold = 0.6) in each cluster were assigned a corresponding letter, resulting in 8 categories: M, T, E, MT, TE, ME, MTE and None (*SI Appendix* **Dataset S6**).

### Statistical analysis

Statistical analyses were performed using GraphPad Prism 8 and R. Data are presented as mean ± standard error of the mean (s.e.m.). An unpaired two-tailed Student’s t-test was used when comparing two groups; an ordinary one-way ANOVA was used when comparing three or more groups. For multiple RCD pathway analysis, the Wilcoxon rank sum test with continuity correction was used. Statistical significance was considered with a P value less than 0.05. \**P* < 0.05, \*\**P* < 0.01, \*\*\**P* < 0.001, and \*\*\*\**P* < 0.0001. All data are presented from at least 3 independent experiments except for snRNA-seq (n = 2/time point).

## Acknowledgements and Funding Sources

## Acknowledgements

This work was performed with the support of the University of Missouri Genomics Technology Core.

## Author Contributions

study conceptualization (ZW, ALP), experimental design (ZW, KMD, SKB, ALP), research conducted (ZW), data analysis (ZW, KMD, SKB, ALP), manuscript preparation (ZW, ALP), manuscript editing (ZW, KMD, SKB, ALP)

