## Supplemental Figs. S1-S5 for "Mesenchymal-epithelial transition serves to rapidly, yet transiently, restore the endometrial epithelium during postpartum murine uterine regeneration"

**Figure S1. Lineage-tracing mouse model. (A)** Schematic of the *Pdgfrα*$^{CreERT2/+}$*; Rosa26-tdTomato*$^{fl/+}$ mouse model. Briefly, the *CreERT2* fusion gene was inserted downstream of the *Pdgfrα* promoter. The *tdTomato* (*tdTom*) gene was inserted downstream of a loxP-floxed transcription terminator sequence (STOP) in the ubiquitous *Rosa26* locus. Upon tamoxifen (TAM) treatment, CreERT2 will translocate to the nucleus and excise the loxP-floxed STOP sequence, resulting in constitutive *tdTom* reporter gene expression. **(B)** Schematic of the experimental design. TAM (2mg/ml) was administered to sexually mature female mice for three consecutive days. Following a 28-day washout, one group of mice were sacrificed at the 1st estrus or diestrus and another group of mice were mated and sacrificed at embryonic (E) 18.5, postpartum day (PPD) 1, 2, 3, 4, and 28. **(C)** Flow cytometry gating strategy of endometrial cells isolated from a littermate control mouse negative for tdTom and stained with EpCAM to establish appropriate gates. Colored squares in the hierarchy plot correspond to populations in the scatter plots. Population counts for a representative mouse are presented in the graph. **(D)** tdTom (red) expression in uteri of *Pdgfrα*$^{CreERT2/+}$*; Rosa26-tdTomato*$^{fl/+}$ mice at estrus and diestrus without and with TAM treatment. Sections were counterstained with DAPI in grey (Merge). **(E)** Quantification of flow cytometry analysis of EpCAM$^{-}$tdTom$^{+}$ stromal cells from mice without and with TAM treatment showing stromal labeling efficiency. **(F)** Quantification of flow cytometry analysis of EpCAM$^{+}$tdTom$^{+}$ epithelial cells without and with TAM in estrus and diestrus. **(G)** Immunofluorescent staining for native Pdgfrα protein (green), with endogenous tdTom expression and nuclear counterstaining with DAPI (grey, Merge). FSC-A: forward scatter area, FSC-H: forward scatter height, SSC-A: side scatter area. Scale bar in all images: 100 µm.

**Figure S2. Quality control of the snRNA-seq data.** Violin plots showing **(A)** number of reads, **(B)** number of genes, **(C)** percentage of mitochondrial genes, and **(D)** percentage of ribosomal genes in nuclei from th indicated mouse uterine samples.

**Figure S3. Regulated cell death pathways in mesenchymal-derived (MD) epithelial cells. (A)** Total, non-MD and MD epithelial cell proportions across postpartum regeneration in *Pdgfrα*$^{CreERT2/+}$*; Rosa26-tdTomato*$^{fl/+}$ mice shows an overall decline to PPD1 with a transient increase at PPD2 primarily due to MD epithelial cells, followed by another decline to PPD3 and another increase at PPD4 primarily due to non-MD epithelial cells. Dot plots of **(B and C)** pro-anoikis genes, **(D and E)** pro-pyroptosis genes, **(F and G)** pro-ferroptosis genes, and **(H and I)** pro-autophagy genes in indicated cell clusters as an aggregate of all time points **(B, D, F and H)** or in non-MD and MD epithelial cell clusters across regeneration time points **(C, E, G, and I)**. Box plots showing the module scores for **(J)** pro-anoikis, **(K)** pro-pyroptosis, **(L)** pro-ferroptosis, and **(M)** pro-autophagy genes in MD and non-MD epithelial cells regeneration time points. A red and blue trend line connecting the mean of the module score was created for MD and non-MD epithelial cells, respectively. * $P < 0.05$, ** $P < 0.01$, *** $P < 0.001$, **** $P < 0.0001$; ns, not statistically different.

**Figure S4. Characterization of transitional cells in PPD1 uteri. (A)** Immunofluorescent (IF) staining of the uterus from a *Pdgfrα*$^{CreERT2/+}$*; Rosa26-tdTomato*$^{fl/+}$ mouse for KRT8 (green) and RFP (magenta) with endogenous tdTom (red) and nuclear counterstaining with DAPI (grey, Merge). Dashed yellow box highlights a region of interest in the stroma (Str) and the solid yellow box is the zoomed-in image showing co-expression of KRT8 and RFP. LE: luminal epithelium. **(B)** IF staining for KRT8 (green) and CD45 (magenta) with nuclear counterstaining with DAPI (grey, Merge) in the uterus. Dashed yellow box highlights a region of interest in the stroma (Str) and the solid yellow box is the zoomed-in image showing a lack of co-expression of KRT8 and CD45. **(C)** IF staining for KRT8 (green) and EpCAM (magenta) with nuclear counterstaining with DAPI (grey, Merge) in the uterus. The KRT8$^+$ transitional cells in the stroma (Str) were negative for EpCAM. Scale bar: 100 μm.

**Figure S5. Inference of the stem/progenitor nature of mesenchymal-derived (MD) transitional cells. (A)** Dotplot of putative endometrial mesenchymal stem cell (eMSC) and endometrial epithelial stem cell (eESC) marker genes in the indicated cell types from *Pdgfrα*$^{CreERT2/+}$*; Rosa26-tdTomato*$^{fl/+}$ mice uteri (aggregate of all time points). Red box highlights MD transitional cluster (MD trans) and dashed red line demarcates gene sets. Violin plots of the module scores of **(B)** eESC and **(C)** eMSC marker genes from (A) in the indicated cell types. Violin plots of **(D)** the module scores of 38 revival stem cell (revSC) signature genes, and **(E)** the module scores of 96 atrophy-induced villus epithelial cell (aVEC) signature genes in the indicated cell types.

### Figure S1

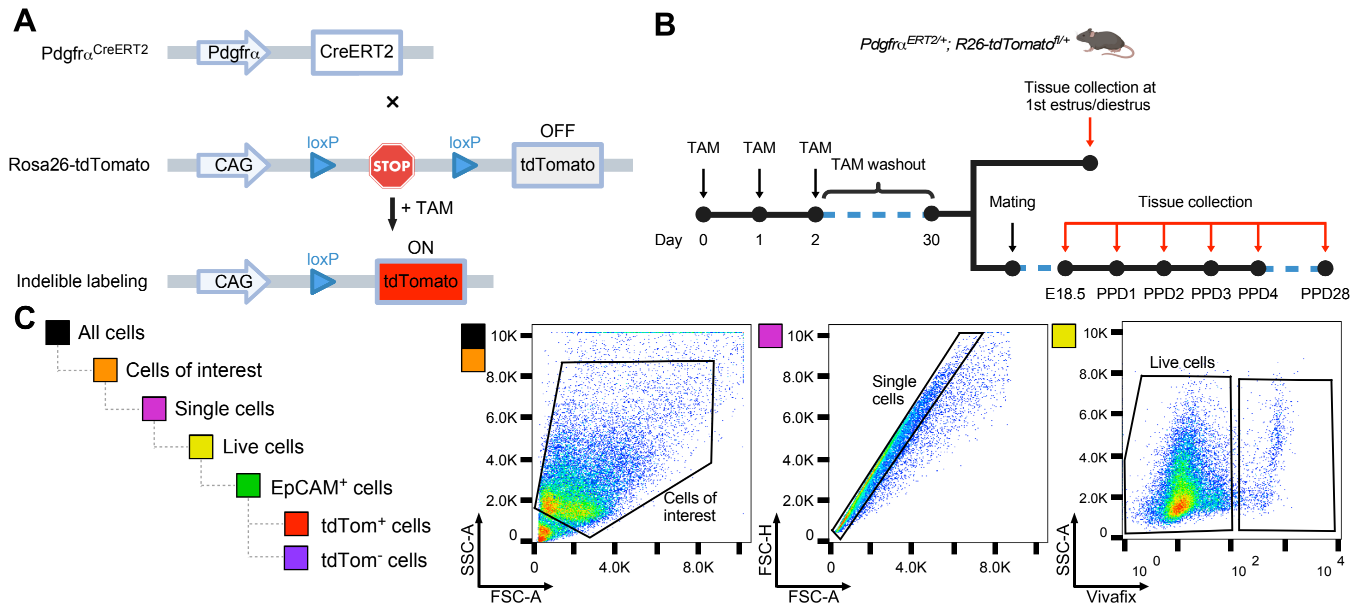


| Cell type | Count | % of Total | % of Plot |
|---|---|---|---|
| All cells | 33,925 | 100 | 100 |
| Cells of interest | 26,910 | 79.3 | 79.3 |
| Single cells | 24,762 | 73.0 | 92.0 |
| Live cells | 23,482 | 69.2 | 94.8 |
| EpCAM+ cells | 4,251 | 12.5 | 18.1 |
| tdTom+ cells | 16 | 0.05 | 0.38 |
| tdTom- cells | 4,210 | 12.4 | 99.0 |

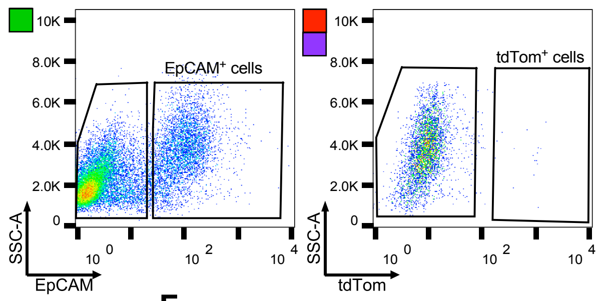


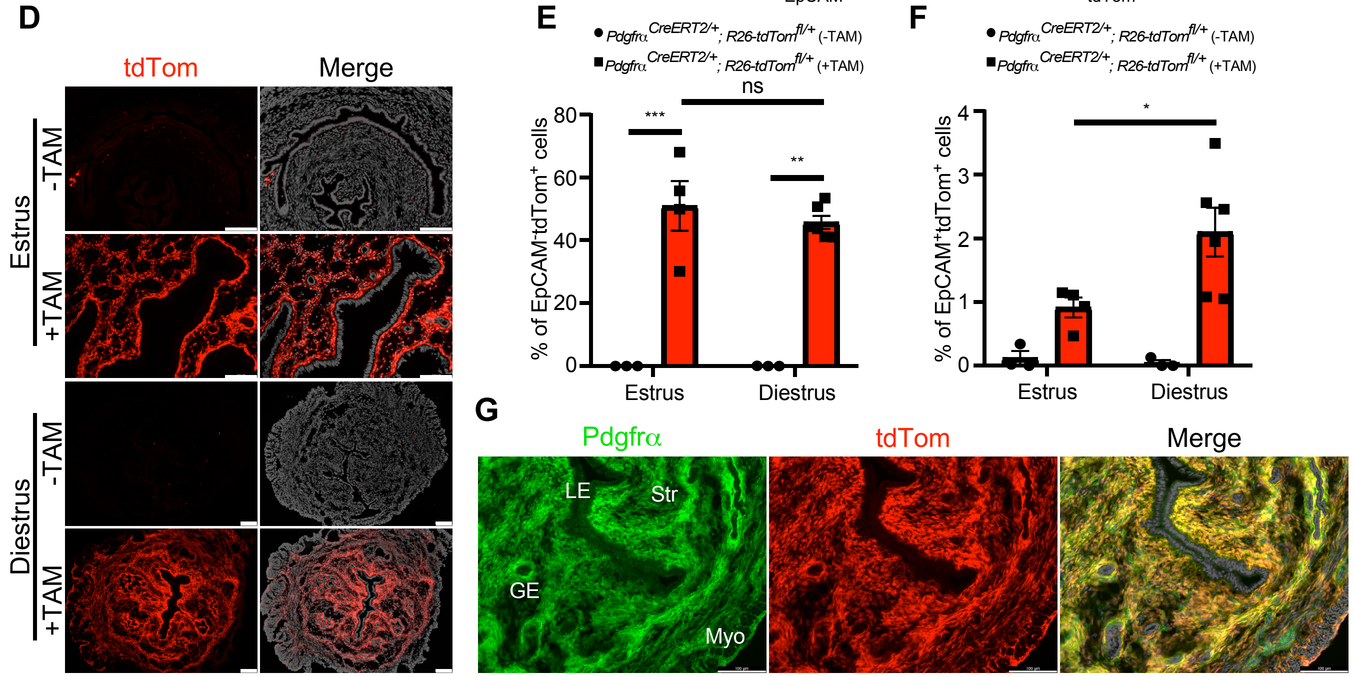

### Figure S2

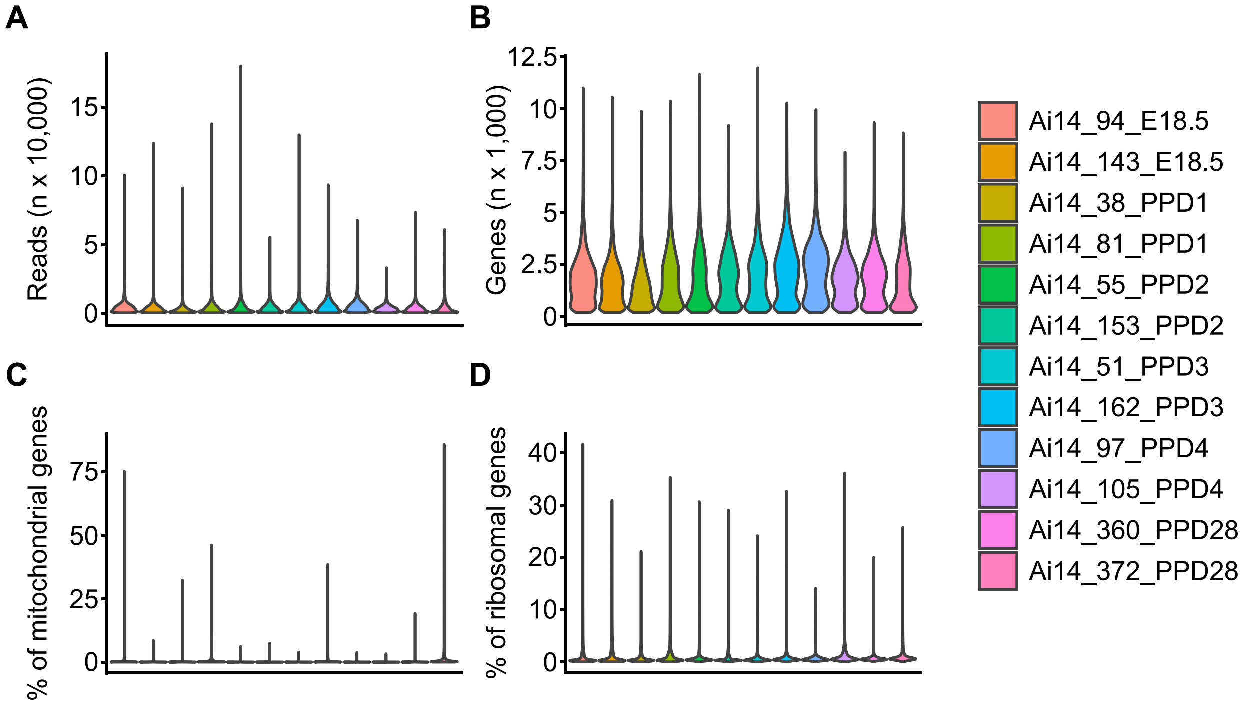
A
B
C
D
Reads (n x 10,000)
0
5
10
15
Genes (n x 1,000)
0
2.5
5
7.5
10
12.5
% of mitochondrial genes
0
25
50
75
% of ribosomal genes
0
10
20
30
40
Ai14_94_E18.5
Ai14_143_E18.5
Ai14_38_PPD1
Ai14_81_PPD1
Ai14_55_PPD2
Ai14_153_PPD2
Ai14_51_PPD3
Ai14_162_PPD3
Ai14_97_PPD4
Ai14_105_PPD4
Ai14_360_PPD28
Ai14_372_PPD28

### Figure S3

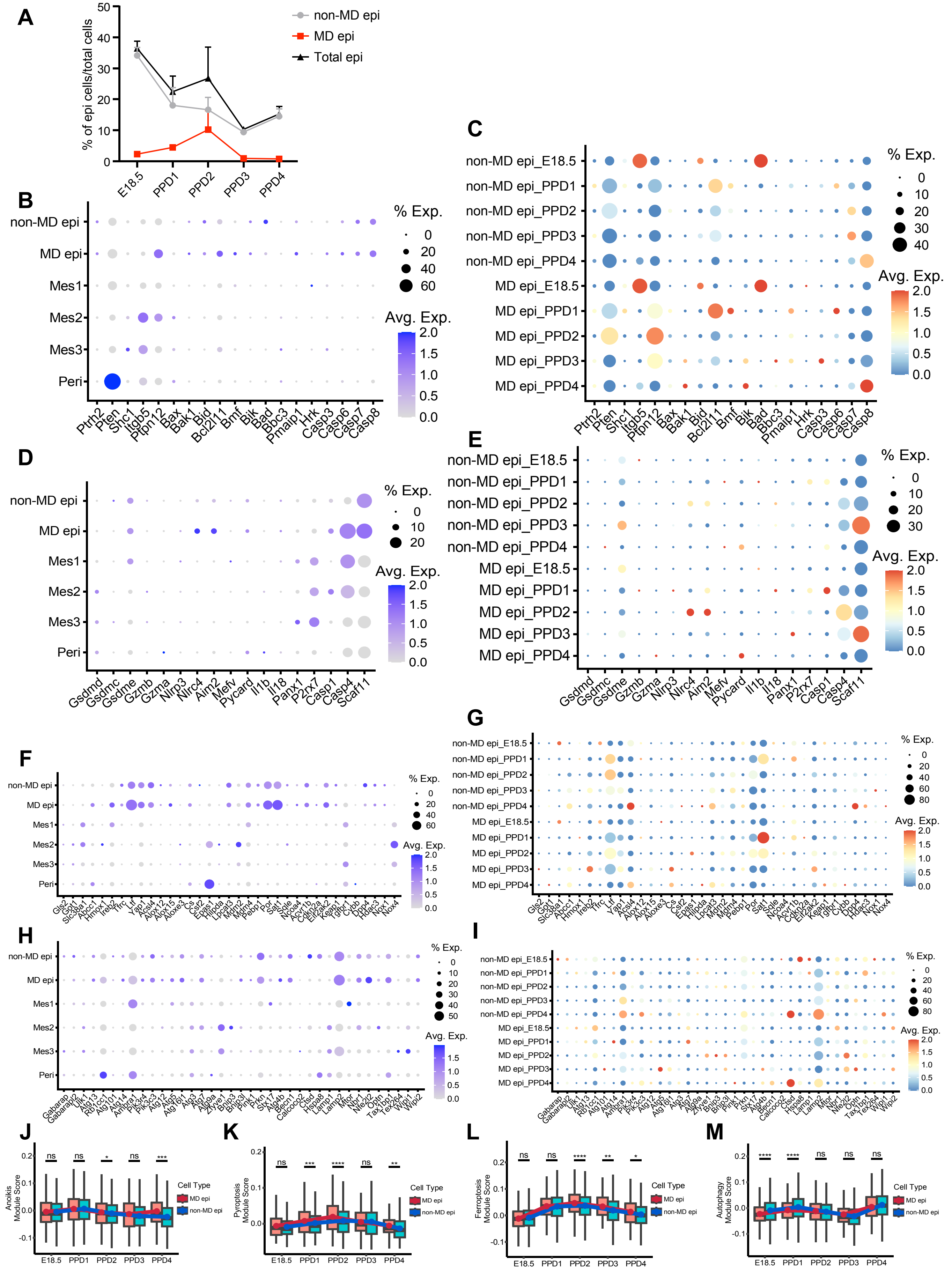
A
% of epi cells/total cells
non-MD epi
MD epi
Total epi
E18.5
PPD1
PPD2
PPD3
PPD4
B
non-MD epi
MD epi
Mes1
Mes2
Mes3
Peri
% Exp.
Avg. Exp.
Ptrh2 Pten Shc1 Itgb5 Ptpn12 Bax Bak1 Bid Bcl2l11 Bmf Bik Bad Bbc3 Pmaip1 Hrk Casp3 Casp6 Casp7 Casp8
C
non-MD epi_E18.5
non-MD epi_PPD1
non-MD epi_PPD2
non-MD epi_PPD3
non-MD epi_PPD4
MD epi_E18.5
MD epi_PPD1
MD epi_PPD2
MD epi_PPD3
MD epi_PPD4
% Exp.
Avg. Exp.
D
Gsdmd Gsdmc Gsdme Gzmb Gzma Nlrp3 Nlrc4 Aim2 Mefv Pycard Il1b Il18 Panx1 P2rx7 Casp1 Casp4 Scaf11
E
F
G
H
I
J
Anoikis Module Score
Cell Type
MD epi
non-MD epi
K
Pyroptosis Module Score
L
Ferroptosis Module Score
M
Autophagy Module Score

### Figure S4

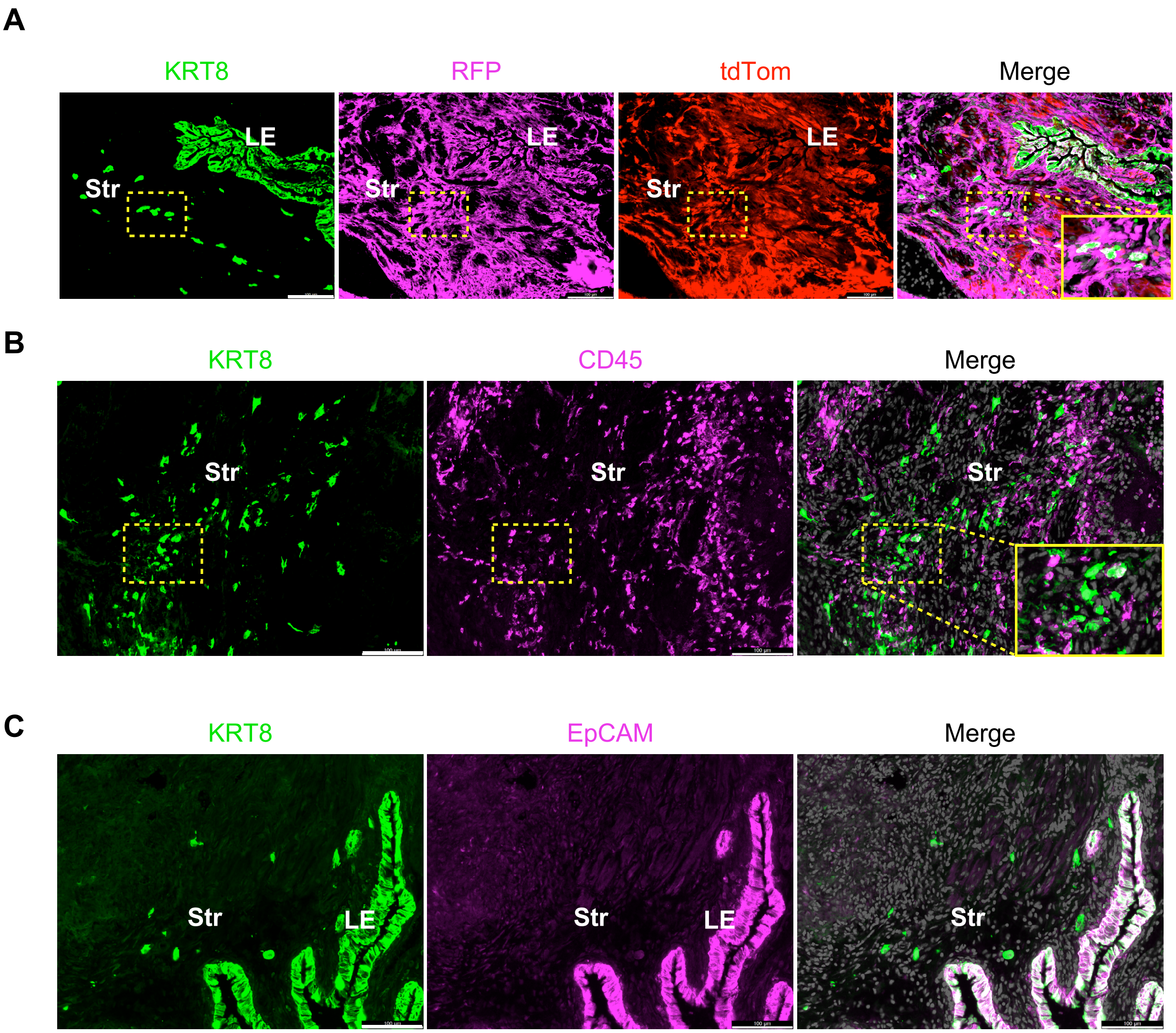
A
KRT8
RFP
tdTom
Merge
LE
Str
B
KRT8
CD45
Merge
Str
C
KRT8
EpCAM
Merge
Str
LE

### Figure S5

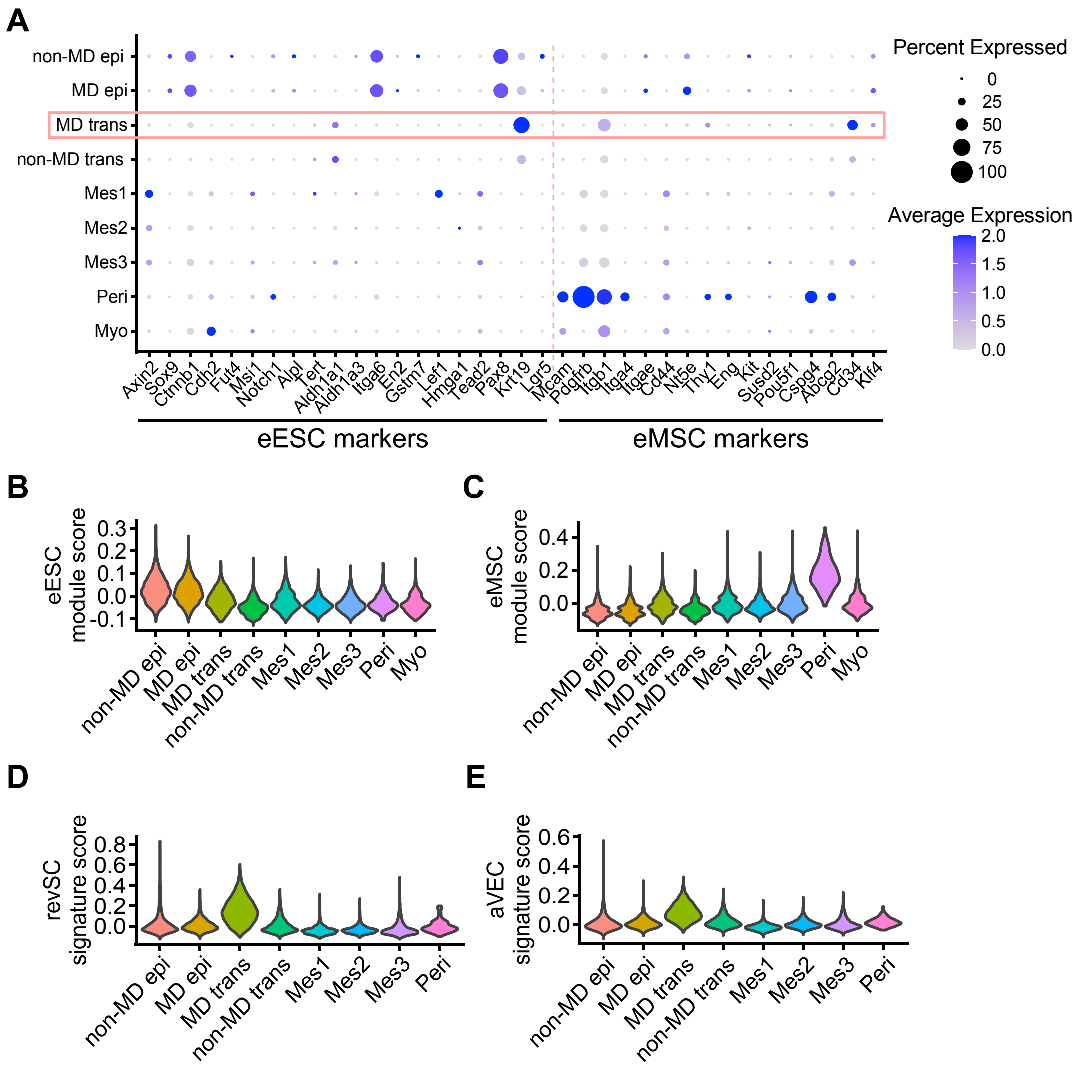

A
non-MD epi
MD epi
MD trans
non-MD trans
Mes1
Mes2
Mes3
Peri
Myo
Axin2
Sox9
Ctnnb1
Cdh2
Fut4
Msi1
Notch1
Alpl
Tert
Aldh1a1
Aldh1a3
Itga6
En2
Gstm7
Lef1
Hmga1
Tead2
Pax8
Krt19
Lgr5
Mcam
Pdgfrb
Itgb1
Itga4
Itgae
Cd44
Nt5e
Thy1
Eng
Kit
Susd2
Pou5f1
Cspg4
Abcg2
Cd34
Klf4
eESC markers
eMSC markers
Percent Expressed
0
25
50
75
100
Average Expression
2.0
1.5
1.0
0.5
0.0
B
eESC module score
C
eMSC module score
D
revSC signature score
E
aVEC signature score
